# MYSM1 co-activates ERα action via histone and non-histone deubiquitination to confer antiestrogen resistance in breast cancer

**DOI:** 10.1101/2022.12.23.521780

**Authors:** Ruina Luan, Ge Sun, Baosheng Zhou, Manlin Wang, Yu Bai, Chunyu Wang, Shengli Wang, Kai Zeng, Jianwei Feng, Mingcong He, Lin Lin, Yuntao Wei, Qiang Zhang, Yue Zhao

## Abstract

Endocrine resistance is a crucial challenge in estrogen receptor alpha (ERα)-positive breast cancer (BCa) therapy. Aberrant alteration in modulation of E2/ERα signaling pathway has emerged as the putative contributor for endocrine resistance in BCa. Thus, identification the efficient ERα cofactor remains necessary for finding a potential therapeutic target for endocrine resistance. Herein, we have demonstrated that Myb like, SWIRM and MPN domains 1 (MYSM1) as a histone deubiquitinase is a novel ERα co-activator with established *Drosophila* experimental model. Our results showed that MYSM1 participated in up-regulation of ERα action via histone and non-histone deubiquitination. We provided the evidence to show that MYSM1 was involved in maintenance of ERα stability via ERα deubiquitination. Furthermore, silencing MYSM1 induced enhancement of histone H2A ubiquitination as well as reduction of histone H3K4me3 and H3Ac levels at *cis* regulatory elements on promoter of ERα-regulated gene. In addition, MYSM1 depletion attenuated cell proliferation/growth in BCa-derived cell lines and xenograft models. Knockdown of MYSM1 increased the sensitivity of antiestrogen agents in BCa cells. MYSM1 was highly expressed in clinical BCa samples, especially in aromatase inhibitor (AI) non-responsive tissues. These findings clarify the molecular mechanism of MYSM1 as an epigenetic modifier in regulation of ERα action and provide a potential therapeutic target for endocrine resistance in BCa.

## Introduction

Breast cancer has been ranked as the malignancy with the highest incidence worldwide since 2020 and owns the highest mortality among cancers in women, which directly impacts their life quality and expectancy (Sung et al, 2021). 70-80% breast cancer is characterized by ERα positive expression (Waks & Winer, 2019). ERα participates in vital cellular processes, such as proliferation, differentiation, and apoptosis of mammary epithelial cells. Given the pleiotropic functions of ERα, perturbation in the estrogen (17β-estradiol, E2)/ERα signaling pathway could result in BCa initiation and progression. The causal role of ERα in BCa pathology makes it a predictive factor and therapeutic target of BCa. At present, endocrine therapy blocking ERα pathway is an exactly prevailing treatment for ERα-positive BCa (Mehta et al, 2019). While most cases are originally sensitive to antiestrogen therapies, the adaptability of tumor cells leads to a substantial percentage of patients stopping responding and gradually developing drug resistance (Hanker et al, 2020). Thus, well understanding the mechanism underlying the modulation of E2/ERα signaling pathway would provide the potential strategies for endocrine resistance.

ERα belongs to a member of the steroid hormone receptor superfamily. E2 binding activates ERα by changing its conformation, thereby transferring into the nucleus to induce the transcription of ERα target genes (Hewitt & Korach, 2018; Yasar et al, 2017). Basic transcriptional machinery along with co-regulators are rapidly recruited to modulate the expression of target genes, such as *c-Myc, CCND1, GREB1, TFF1* (Metivier et al, 2003; Shang et al, 2000). The complicated network of these co-regulators defines a code, which acts as an adapter molecule connecting ERα to the basal transcription apparatus or alters chromosome structure on cis-regulatory elements, comprising covalent modifications in histone tails and nucleosome remodeling (Dimitrakopoulos et al, 2021). Accumulating evidence suggests that the core regulatory proteins of ERα action subtly tune hormone sensitivity, receptor stability, and ERα-mediated transcriptional activity according to their diverse enzymatic activities and associated patterns (Manavathi et al, 2013; Shao et al, 2004; Sukocheva et al, 2020). A series of ERα cofactors have been identified so far to regulate the estrogen-driven transcriptional program. CBP/p300, p/CAF, the p160 family, PELP1, SWI/SNF complex, YAP, and PARP-1 identified as ERα co-activators lead to chromatin de-condensation and modulate epigenetic changes, providing a selective advantage for cancer cell growth, differentiation, invasion, metastasis, and endocrine resistance (Gadad et al, 2021; Garcia-Martinez et al, 2021; Ju et al, 2006; Schiewer & Knudsen, 2014; Zhu et al, 2019). While the co-repressors of ERα, such as SMRT, NCOR1, PIP140, BRCA1, MTA1, TLE3, play multiple functions on breast cancer processes through modulation of ERα action (Dobrzycka et al, 2003; Wen et al, 2009). It is convictive that accurate orchestration of ERα action results from the coordination of multiple co-regulators. Since ER conducts a diverse function, identification of novel ERα co-regulator is still necessary for finding the potential target for ERα-positive breast cancer treatment.

MYSM1 is a metalloproteinase with deubiquitinase catalytic activity. It enhances chromatin accessibility by specifically removing H2Aub to facilitate gene transcription (Zhu et al, 2007). MYSM1 is involved in extensive physiological processes. In the hematopoietic system, MYSM1 de-represses an array of genes which are pivotal for lineage specification and stem cell differentiation through histone H2A deubiquitination (Belle et al, 2020; Nijnik et al, 2012; Wang et al, 2013). Additionally, MYSM1 guards against excessive inflammation and autoimmune reaction under circumstances of inflammatory irritation or infection by non-histone deubiquitination to abrogate NOD2:RIP2, cGAS-STING, or TRAF3/6 signaling in the immune system (Panda & Gekara, 2018; Panda et al, 2015; Tian et al, 2020b). MYSM1 deficiency spontaneously perturbs the proper repair of DNA damage to accumulate DNA double strand breaks (DSB), accompanied by cellular senescence (Kroeger et al, 2020; Nishi et al, 2014; Tian et al, 2020a). MYSM1 is also involved in tumor pathologic processes. It has been reported that MYSM1 suppresses colorectal cancer progression via histone H2A deubiquitination, thereby activating miR-200 family members/CDH1 (Chen et al, 2021). MYSM1 co-activates androgen receptor (AR) action to promote prostate cancer (Zhu et al, 2007), while MYSM1 inhibits castration-resistant prostate cancer (CRPC) process through PI3K/Akt signaling regulation (Sun et al, 2019). In triple-negative breast cancer (TNBC), MYSM1 reduces RSK3 protein to inactive RSK3-phospho-BAD pathway and induces apoptosis and cisplatin sensitivity (Guan et al, 2022). However, the molecular mechanisms underlying the modulation function of MYSM1 on ERα signaling in ERα-positive BCa remains elusive.

In this study, we identify MYSM1 as a co-activator of ERα in a *Drosophila* model carrying an ERα-mediated gene transcription system. We further demonstrate that MYSM1 associates with ERα and increases ERα-induced transcriptional activity in breast cancer cell lines. Knockdown of MYSM1 inhibits transcription of the endogenous ERα target genes. MYSM1 is recruited together with ERα to the promoter region of E2-induced genes, leading to epigenetic modulation of H2Aub, H3K4me3, and H3Ac levels. Unexpectedly, our data suggests that MYSM1 stabilizes the ERα protein on Lysine 48 (K48) and K63-linked poly-ubiquitination via its deubiquitinase activity. MYSM1 depletion suppresses cell proliferation and increases antiestrogen sensitivity in BCa. Furthermore, MYSM1 is highly expressed in clinical BCa samples, and higher expression of MYSM1 predicts a poor survival of BCa patients. Taken together, our findings suggest that the deubiquitinase activity of MYSM1, in addition to its epigenetic modulation to function as a co-activator of ERα, is involved in non-histone modification to maintain ERα stability in ERα-positive BCa progression.

## Results

### MYSM1 physically associates with ERα in breast cancer cells

We have previously identified several co-regulators of AR in *Drosophila* experimental system (Sun et al, 2016; Wang et al, 2015). Herein, we generated an inducible *Drosophila* model containing an ERα-mediated gene transcription system to isolate co-regulators involved in modulating ERα actions. The system includes a GAL4 expression construct, a UAS-linked ERα expression construct, an ERE-linked green fluorescent protein (GFP) reporter construct with the modified GAL4-UAS bipartite approach in *Drosophila* (Fig 1A). ERα-induced transactivation can be monitored by the intensity of GFP expression in *Drosophila* experimental system. The experimental *drosophila* model was crossed with *Drosophila* mutants of potential genes, and GFP protein expression was assessed to screen ERα cofactors in the presence of E2. Interestingly, loss of function mutant of *CG4751* dramatically compromised ERα-induced GFP protein expression. To confirm this result, we further generated transgenic *Drosophila* with human homologue of *CG4751* (*MYSM1*), the results demonstrated that MYSM1 increased ERα-mediated transactivation, suggesting that MYSM1 may be a co-activator of ERα (Fig EV1A).

**Figure 1.**
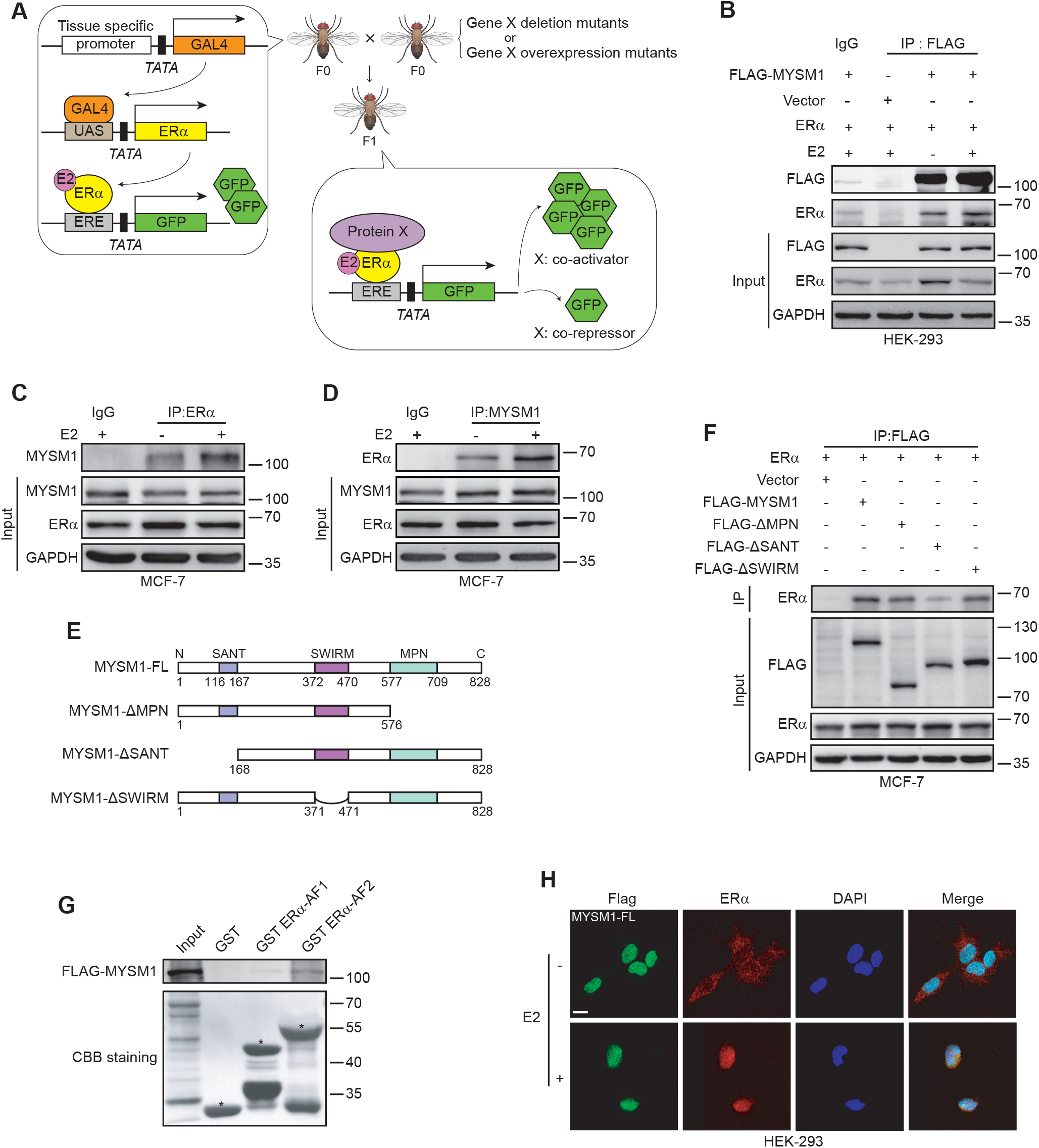
MYSM1 interacts with ERα in breast cancer cells. A. Co-immunoprecipitation showing the interaction between exogenous MYSM1 and ERα with or without E2. HEK293 cells were co-transfected with FLAG-tagged MYSM1 and ERα plasmids and incubated in estrogen-depleted medium (5% charcoal-stripped serum in phenol red-free DMEM) for 48 h, then treated with ethanol vehicle or E2 (100 nM) for 12 h. Whole-cell extracts were harvested for immunoprecipitation with the anti-FLAG or IgG antibodies, with 5% reserved as the input control. B, C. Reciprocal Co-immunoprecipitation to detect the association between endogenous MYSM1 and ERα in MCF-7 cells with the indicated antibodies. D. Schematic diagram representing the MYSM1 full length and truncated expression plasmids (ΔMPN, ΔSANT, ΔSWIRM) containing FLAG tags. E. Co-immunoprecipitation to determine the exact domains in MYSM1 responsible for its binding to ERα. Ectopic ERα along with wild-type MYSM1 or MYSM1 deletion mutants were expressed in MCF-7 cells. Anti-FLAG-MYSM1 immunoprecipitates or corresponding input were then immunoblotted for ERα, MYSM1 (FLAG) and GAPDH. F. Detection of the directly binding of MYSM1 and ERα by GST-pull down assay. GST, GST ERα-AF1, GST ERα-AF2 proteins expressed in an *E. coli* system were incubated with the *in vitro* transcribed 35S-MYSM1 protein. The binding proteins in the reaction mixtures were analyzed by SDS-PAGE and autoradiography. Asterisks marked GST fusion protein locations. G. Immunofluorescent staining of MYSM1 (anti-FLAG, green) and ERα (anti-ERα, red) in HEK293 cells overexpressing MYSM1 truncated mutants and ERα in the presence of E2. Nuclei were stained with DAPI (blue). Scale bars, 10 µm.

We thus turn to ask whether MYSM1 associates with ERα. STRING database (http://www.bork.embl-heidelberg.de/STRING/) predicted that MYSM1 may correlate with ERα (Fig EV1B). Then, we set out to examine the interaction between MYSM1 and ERα in mammalian cells by Co-immunoprecipitation (Co-IP). ERα and FLAG-tagged MYSM1 expression plasmids were co-transfected into HEK293 cells. And specific antibodies against FLAG was used to precipitate the related protein as indicated. Our results showed that MYSM1 associated with ERα (Fig 1B). The similar experiments were performed in ERα-positive breast cancer (BCa)-derived cell lines (MCF-7 and T47D). The results showed that the endogenous MYSM1 associated with ERα (Fig 1C and D, and Fig EV1C and D). To determine the exact interaction region in MYSM1, we constructed MYSM1 truncated mutant plasmids as indicated (Fig 1E). Co-IP experiments were performed with co-transfection of these plasmids and ERα expression plasmid in mammalian cells. The precipitation results demonstrated that loss of SANT (Swi3, Ada2, N-CoR, (TF) IIIB) domain more obviously impaired the association between MYSM1 and ERα, compared with that of SWIRM or MPN domain (Fig 1F, and Fig EV1E). Additionally, GST-pull down was performed with GST-tagged ERα-AF1 or ERα-AF2 fragments. The results showed that MYSM1 directly interacted with ERα-AF2 fragment (Fig 1G).

To investigate the subcellular distribution of MYSM1 and ERα, we applied immunofluorescence (IF) in HEK293 cells and breast cancer cells (MCF-7 and T47D) (Fig 1H, Fig EV1F-H). We observed that MYSM1 is distributed in the nucleus regardless of E2 (10^−7^ M) treatment, whereas ERα diffused into the nucleus and entirely co-located with MYSM1 in the presence of E2. HEK293 cells were transfected with MYSM1 truncated mutant plasmids and ERα expression plasmid. The results from IF experiments demonstrated that three kinds of mutants of MYSM1 (deletion of MPN, SWIRM, or SANT) were distributed in the nucleus, indicating that nuclear localization signal (NLS) of MYSM1 may exist in other regions except MPN, SWIRM, and SANT domains (Fig EV1H). Collectively, these data suggest that MYSM1 interacts with ERα.

### MYSM1 co-activates ERα-induced transcriptional activity

Having established that ERα-mediated transactivation is significantly down-regulated by loss of function of CG4751 (*Drosophila* homologue of MYSM1) in *Drosophila* experimental system, we speculated that MYSM1 may functionally participates in ERα signaling pathway in human. Luciferase assays was then performed to examine the effect of MYSM1 on modulation of ERα action. The results demonstrated that MYSM1 co-activated ERα-induced transactivation in a dose-dependent manner in HEK293 cells (Fig 2A, and Fig EV2A). Meanwhile, MYSM1 knockdown inhibits ERα action in T47D cells, indicating that MYSM1 is a novel co-activator of ERα (Fig Fig EV2C). ERα mainly contains N-terminal activation function-1 (ERα AF-1) carrying constitutive ligand-independent domain, and C-terminal activation function-2 (ERα AF-2) harboring ligand-dependent domain. We sought to examine the effect of MYSM1 on transactivation induced by the truncated ERα. The results showed that MYSM1 simultaneously increased ERα AF-1 and AF-2 actions (Fig 2B, and Fig EV2B). To gain insight into the functional domain of MYSM1, we further performed the similar experiments with MYSM1 truncated mutant plasmids. Compared with the full length of MYSM1, the lack of SWIRM domain had little effect on ERα-induced transactivation, while MYSM1-ΔSANT or MYSM1-ΔMPN dramatically reduced ERα action (Fig 2C), indicating that the SANT or MPN domain is required for the co-activation function of MYSM1 on ERα-mediated transactivation.

**Figure 2.**
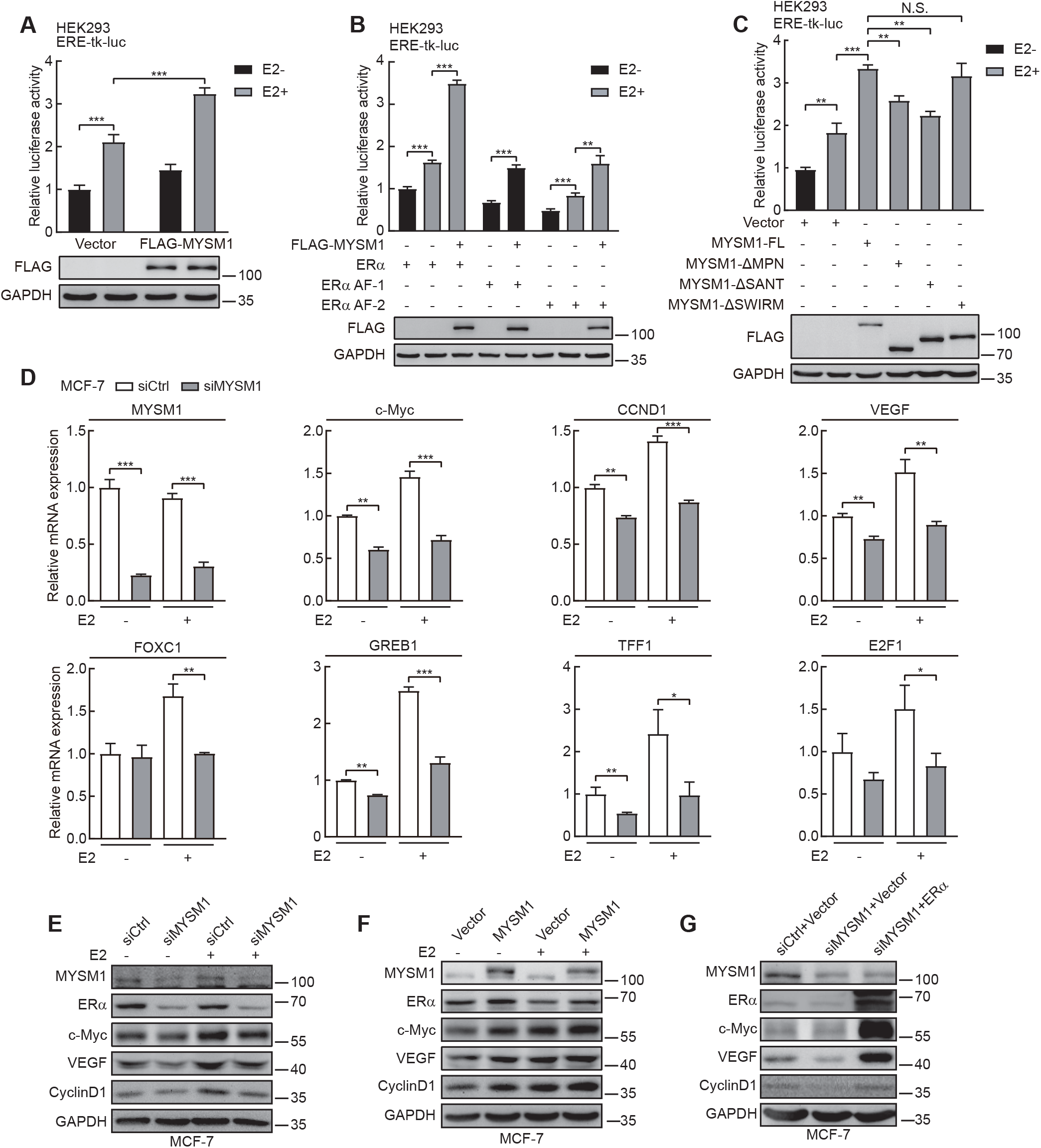
MYSM1 enhances ERα-mediated gene transcription in mammalian cells. A. Relative luciferase activities in HEK293 cells transfected with ERα expression plasmid, ERE-tk-luc, pRL-tk, and the indicated expression plasmids in the presence or absence of E2 (100nM). The expression of MYSM1 was detected by western blot. \*\*\**P* < 0.001 (mean ± SD; Student *t*-test). B. MYSM1 increases ERα AF1 and ERα AF2 mediated transcriptional activity. HEK293 cells were transfected with ERα full length or truncated mutants harboring ERα AF-1 or ERα AF-2 with or without MYSM1 expression. \*\**P* < 0.01, \*\*\**P* < 0.001 (mean ± SD; Student *t*-test). C. The effect of a domain-defective mutation of MYSM1 (ΔMPN, ΔSANT, and ΔSWIRM) on luciferase activity in dual-luciferase reporter system transfected HEK293 cells. The expression of MYSM1 and its truncated mutants were examined with anti-FLAG by western blot. \*\**P* < 0.01, \*\*\**P* < 0.001 and N.S. stands for no significance (mean ± SD; Student *t*-test). D. qPCR analysis to determine the transcript amounts of certified ERα target genes in MCF-7 cells with MYSM1-depleted. \**P* < 0.05; \*\**P* < 0.01; \*\*\**P* < 0.001 (mean ± SD; Student *t*-test). E, F. Immunoblot of ERα target gene expression using the indicated antibodies in MYSM1-depleted MCF-7 cells (E) and MYSM1-overexpressed MCF-7 cells (F). G. The loss of ERα and its target genes in MYSM1-depleted MCF-7 cells can be rescued by ectopic ERα expression. MCF-7 cells were transfected with control siRNA (siCtrl) or siRNA specific against MYSM1 (siMYSM1) followed by PcDNA3.1/ERα expression plasmid.

Indeed, ERα behaves in numerous biological processes through its target genes. To confirm the effect of MYSM1 on regulation of E2/ERα signaling pathway, quantitative real-time PCR (qPCR) experiments were performed to examine co-activation function of MYSM1 on ERα target gene transcription in ERα-positive breast cancer cells. The data demonstrated MYSM1 knockdown significantly suppressed the transcription of endogenous ERα regulated genes, including *c-Myc, CCND1, VEGF, GREB1, TFF1, E2F1, FOXC1* (Fig 2D, and Fig EV2D). Western blot was further conducted for examining the regulation function of MYSM1 on ERα action. The results showed that MYSM1 depletion inhibited the protein levels of ERα regulated genes, such as c-Myc, CCND1, and VEGF (Fig 2E, and Fig EV2E). Consistent with this, ectopic expression of MYSM1 increased c-Myc, CCND1 and VEGF protein level (Fig 2F, and Fig EV2F). Moreover, additional transfection with ERα expression plasmids could reverse the reduced expression of ERα target genes caused by siRNA against MYSM1 (siMYSM1) in breast cancer cells (Fig 2G, and Fig EV2G). Our results suggest that MYSM1 acts as a novel co-activator of ERα.

### MYSM1 stabilizes the ERα protein with its deubiquitination activity

Unexpectedly, in Co-IP experiments as shown in Figure 2E and F, we observed that ectopic expression of MYSM1 increased ERα protein expression, while knockdown of MYSM1 reduced ERα protein level. This prompted us to ask how MYSM1 influences ERα protein expression in breast cancer. shRNA lentivirus against MYSM1 (shMYSM1) was infected into ER-positive breast cancer cell lines. The results showed that knockdown of MYSM1 decreased ERα protein level, whereas ERα mRNA expression had no significant change (Fig 3A and B). On the other hand, ectopic expression of MYSM1enhanced ERα expression in MCF-7 and HEK293 (Fig 3D), indicating that MYSM1 up-regulates ERα protein itself at the post-transcriptional level. When cycloheximide (CHX) was applied at specified time points to inhibit protein synthesis, depletion of MYSM1 significantly accelerated ERα degradation and ectopic expression of wild-type MYSM1 plasmid (MYSM1-FL) ameliorated ERα degradation process (Fig 3C and E). In addition, we treated MCF-7 cells with proteasome inhibitor (MG132), the results showed that regulation of MYSM1 on ERα protein expression was prevented by MG132 treatment (Fig 3F and G), suggesting that MYSM1 is involved in maintenance of ERα protein stability.

**Figure 3.**
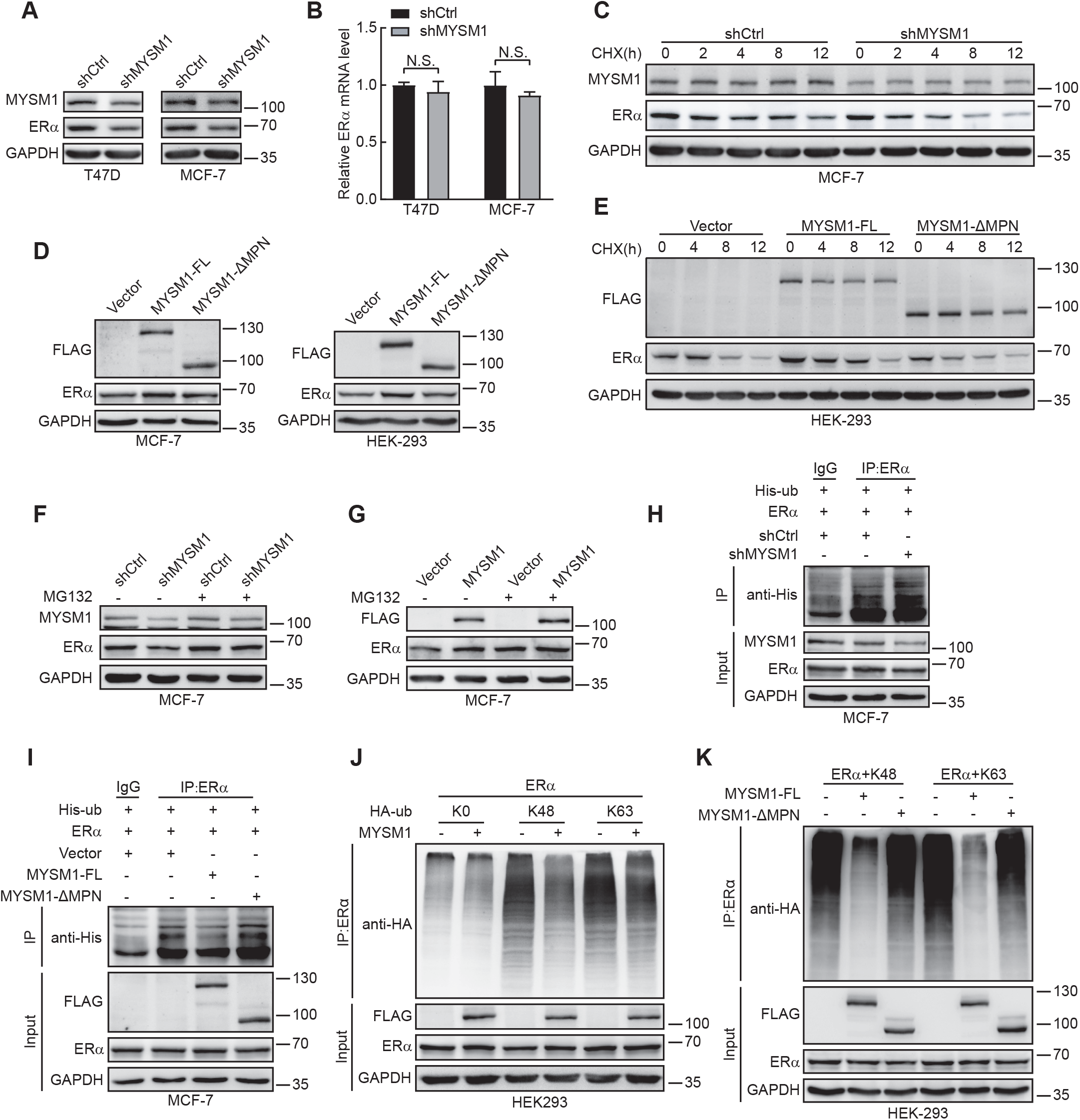
MYSM1 stabilizes the ERα protein through ERα deubiquitination. A, B. Western blot (A) and qPCR (B) analysis in T47D/MCF-7 cells infected with shCtrl or shMYSM1 lentivirus to evaluate the impact of MYSM1 depletion on ERα protein and mRNA levels. N.S. means no significance (mean ± SD; Student *t*-test). C. MCF-7 cells carrying control shRNA or shMYSM1 were complemented with CHX (20 μM) for particular time periods, and cell lysates were assessed by Western blot. D. Western blot analysis of ERα and exogenous MYSM1 expression in MCF-7/HEK293 cells transfected with wild-type MYSM1 plasmid or MPN-deletion mutant. E. Cell lysates from MYSM1-FL/MYSM1-ΔMPN overexpressing MCF-7 cells stimulated with CHX (20 μM) at specified time points were denatured and quantitated by western blot. F, G. Extracts obtained from MCF-7 cells harboring shMYSM1 or FLAG-MYSM1 upon MG132 (10 μM) treatment for 4 h were analyzed by western blot. H. MCF-7 cells from the control or the MYSM1-knockdown group were harvested after MG132 (10 μM) addition for 6h, followed by immunoprecipitation with anti-ERα and subsequently probed with anti-His. I. Immunoblot analysis of the polyubiquitination of ERα proteins in MCF-7 cells transiently co-transfected with plasmids encoding ERα and FLAG-MYSM1 or MYSM1-ΔMPN. MG132 (10 μM) was added to cell culture 6 h before cell collection. Cell lysates were subjected to immunoprecipitation with anti-ERα and immunoblot with anti-His. J, K. HEK293 cells transfected with the indicated plasmids were extracted and immunoprecipitated with anti-ERα, followed by immunoblot with anti-HA.

MYSM1 belongs to the JAMM family of deubiquitinase, so we hypothesized it may participate in maintenance of ERα stability through its deubiquitinase activity. Next, we turned to perform western blot to detect the effect of full length of MYSM1 (MYSM1-FL) and catalytically loss of function of MYSM1 mutant (MYSM1-ΔMPN) on ERα protein expression. The results demonstrated MYSM1-ΔMPN largely decreased the enhancement of ERα expression induced by MYSM1-FL (Fig 3D and E), indicating that deubiquitinase activity of MYSM1 is indispensable for maintenance of ERα stabilization. Ubiquitination assays based on immunoprecipitation were further performed to determine whether ERα is a substrate of MYSM1. Polyubiquitinated ERα proteins were visibly enriched by immunomagnetic beads in the control group, and MYSM1 depletion significantly enhanced ubiquitination of ERα (Fig 3H). In addition, the level of ERα ubiquitination underwent a sharp decline by MYSM1, while seemed to remain constant with MYSM1-ΔMPN, indicating that MYSM1 is involved in ERα deubiquitination (Fig 3I). So far, the most thoroughly characterized polyubiquitin processes are lysine 48 (K48)-and K63-linked ubiquitination. K48-linked chains prefer to target proteins for proteasomal degradation, while K63-linked chains act as a molecular glue for complex formation in various signaling pathways and also participate in protein degradation (Grice & Nathan, 2016; Hayden & Ghosh, 2008; Liu et al, 2018; Madiraju et al, 2022; Wang et al, 2022). We next examined which sites of polyubiquitin chains attached to ERα could be removed by MYSM1. HEK293 cells were transfected with HA-tagged Ubiquitin (Ub) K0, K48, or K63 plasmid for ubiquitination assays. K0 mutant lacking lysine residues was used as a negative control. Ectopic expression of MYSM1 contributes to reduction on K48 and K63 polyubiquitin of ERα (Fig 3J). We then sought to determine the requirement of MYSM1 deubiqitination activity for its ubiquitin linkages cleavage by ubiquitination assay with MYSM1-FL and MYSM1-ΔMPN. Our results demonstrated that catalytically loss of function of MYSM1 mutant (MYSM1-ΔMPN) abrogated the deubiquitination level of K48, K63-linked ubiquitin chains on ERα induced by MYSM1-FL (Fig 3K). Taken together, our data suggest that MYSM1 removes K48 and K63-polyubiquitin conjugates of ERα via its deubiquitination activity, participating in the maintenance of ERα stability.

### MYSM1 is recruited to the promoter region of ERα target gene to be involved in histone H2A deubiquitination

To further assess the epigenetic mechanism underlying the modulation of MYSM1 on ERα action, chromatin immunoprecipitation (ChIP) assay was performed to examine the recruitment of MYSM1 or ERα on estrogen response elements (EREs) on the upstream of the transcription start site (TSS) of c-Myc, which is a putative ERα target gene (Sun et al, 2020) (Fig 4A). The results showed that MYSM1 or ERα was recruited to the ERE region of c-Myc upon E2 treatment (Fig 4B). MYSM1 as a deubiquitinase is involved in deubiquitination of histone H2A to enhance gene transcription. To study the influence of MYSM1 on histone modification on the EREs of ERα target gene, ChIP assays were further performed in BCa cells with siRNA against MYSM1 (siMYSM1). The results demonstrated MYSM1 or ERα is recruited to ERE region on the promoter of *c-Myc* upon E2 treatment. MYSM1 depletion decreased ERα recruitment on EREs (Fig 4C and Fig EV3A). Importantly, MYSM1 knockdown enhanced histone H2Aub level. Moreover, we also observed that MYSM1 depletion inhibited H3K4me3 and H3Ac levels, while H2Bub1 level had no significant change on ERE region of *c-Myc* in BCa-derived cells. To further determine whether MYSM1 and ERα are recruited together to the promoter of *c-Myc*. ChIP-re-ChIP assays were performed. The data showed that MYSM1 and ERα are recruited to ERE region of c-Myc promoter at the same time in the presence of E2 (Fig 4D and E, and Fig EV3B and C). ChIP assays was further conducted to detect the recruitment of MYSM1 or ERα to estrogen response elements of a number of ERα target genes, including *c-Myc, CCND1, VEGF, TFF1* and *GREB1* (Sun et al, 2020). We observed that MYSM1 or ERα was recruited to EREs on these genes, and ERα enrichment at ERE regions of these genes displayed a descended trend upon MYSM1 knockdown with E2 treatment (Fig 4F). Collectively, the data suggest that knockdown of MYSM1 reduces histone H2Aub level on EREs, and declines ERα recruitment to EREs on ERα target genes.

**Figure 4.**
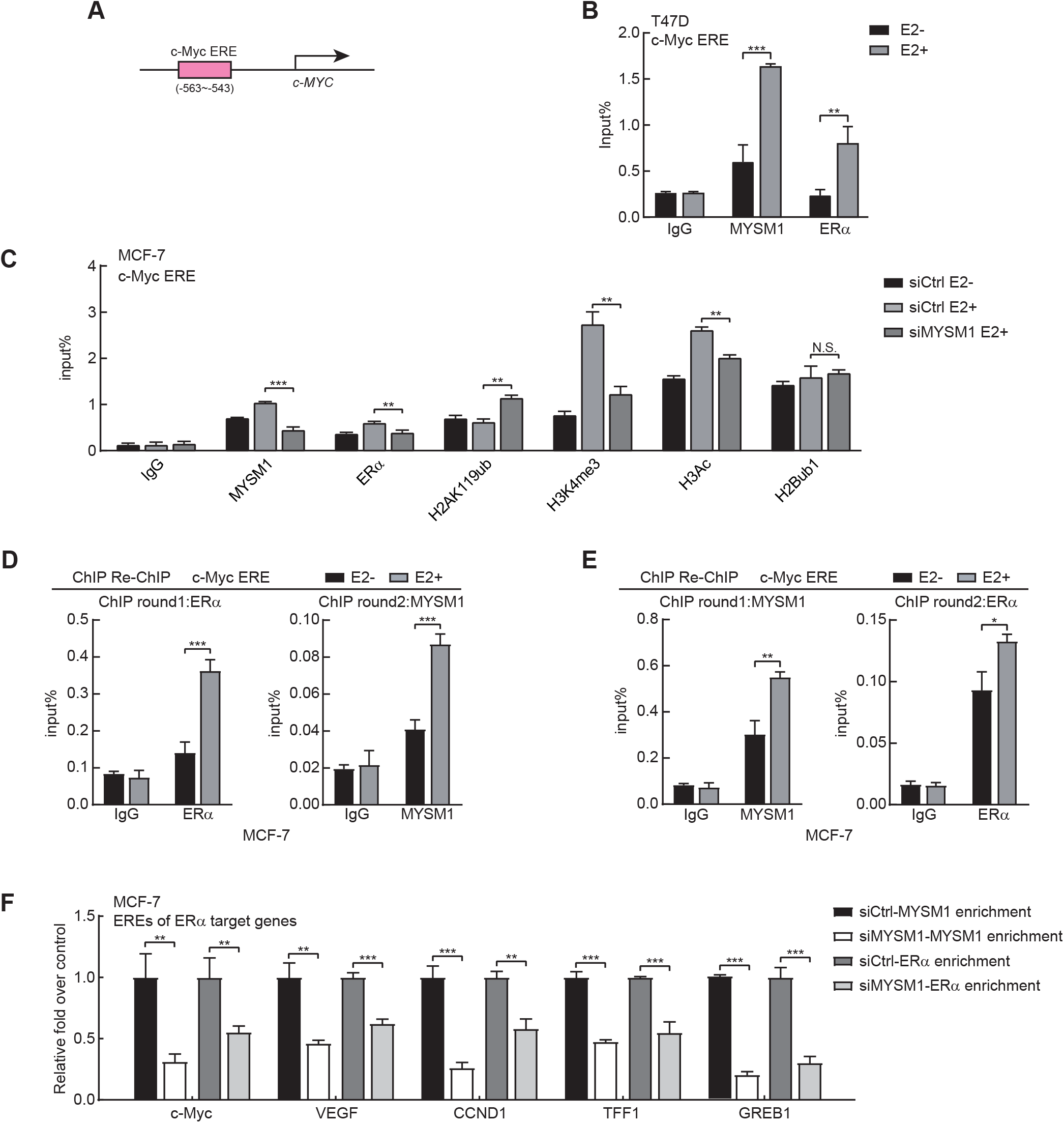
MYSM1 orchestrates ERα occupancy and histone modifications at the locus of ERα target gene. A. Schematic diagram of the putative estrogen response element (ERE) of c-Myc. B. ChIP assay with indicated antibodies shows the recruitment of MYSM1 and ERα at c-Myc ERE. \*\**P* < 0.01; \*\*\**P* < 0.001 (mean ± SD; Student *t*-test). C. ChIP assays via designated antibodies to elaborate the influence of MYSM1 regarding to ERα recruitment and relative histone modifications on *c-Myc* ERE. \*\**P* < 0.01, \*\*\**P* < 0.001 (mean ± SD; Student *t*-test). D, E. ChIP re-ChIP experiments validated that E2 treatment augmented concurrent recruitment of MYSM1 and ERα on *c-Myc*-ERE in MCF-7 cells. Chromatin extracts immunoprecipitated with anti-ERα (D) or anti-MYSM1 (E) as the round1 antibody were re-immunoprecipitated with anti-MYSM1 (D) or anti-ERα (E) respectively, followed by qPCR to calculate the collected DNA signals. \**P* < 0.05; \*\**P* < 0.01; \*\*\**P* < 0.001 (mean ± SD; Student *t*-test). F. MYSM1 and ERα ChIP analysis was conducted in MYSM1-deficiency MCF-7 cells on some ERα-binding sites of E2-induced genes as indicated. Cells were cultured in estrogen-depleted medium for 48 h before stimulation with 100 nM E2 for 2 h. The immunoprecipitated DNA fragments were analyzed by qPCR using primers recognizing the promoter region of ERα target genes. The amplified products were standardized by a certain amount of unprecipitated input DNA. \**P* < 0.05; \*\**P* < 0.01; \*\*\**P* < 0.001 (mean ± SD; Student *t*-test).

### MYSM1 accelerates cell proliferation in ERα-positive breast cancer cell lines

Having established the molecular mechanisms of MYSM1 on ERα action, we then turned to interrogate the potential biological function of MYSM1 on ER-positive breast cancer progression. The results from colony formation assay showed that MYSM1 depletion suppressed colony formation in MCF-7 and T47D cells (Fig 5A and Fig EV4A). Consistently, growth curve analysis showed that knockdown of MYSM1 inhibited cell proliferation, and ectopic expression of ERα partly rescued this condition (Fig 5B and Fig EV4B). We then performed flow cytometry for detecting the influence of MYSM1 on cell cycle as displayed. The results showed that MYSM1 depletion retarded the G1-S phase transition (Fig 5C and Fig EV4C-E). Above all, the results indicate that MYSM1 promotes cell proliferation and G1-S phase transition in BCa, and the effect of MYSM1 on cell growth is partially related to ERα pathway.

**Figure 5.**
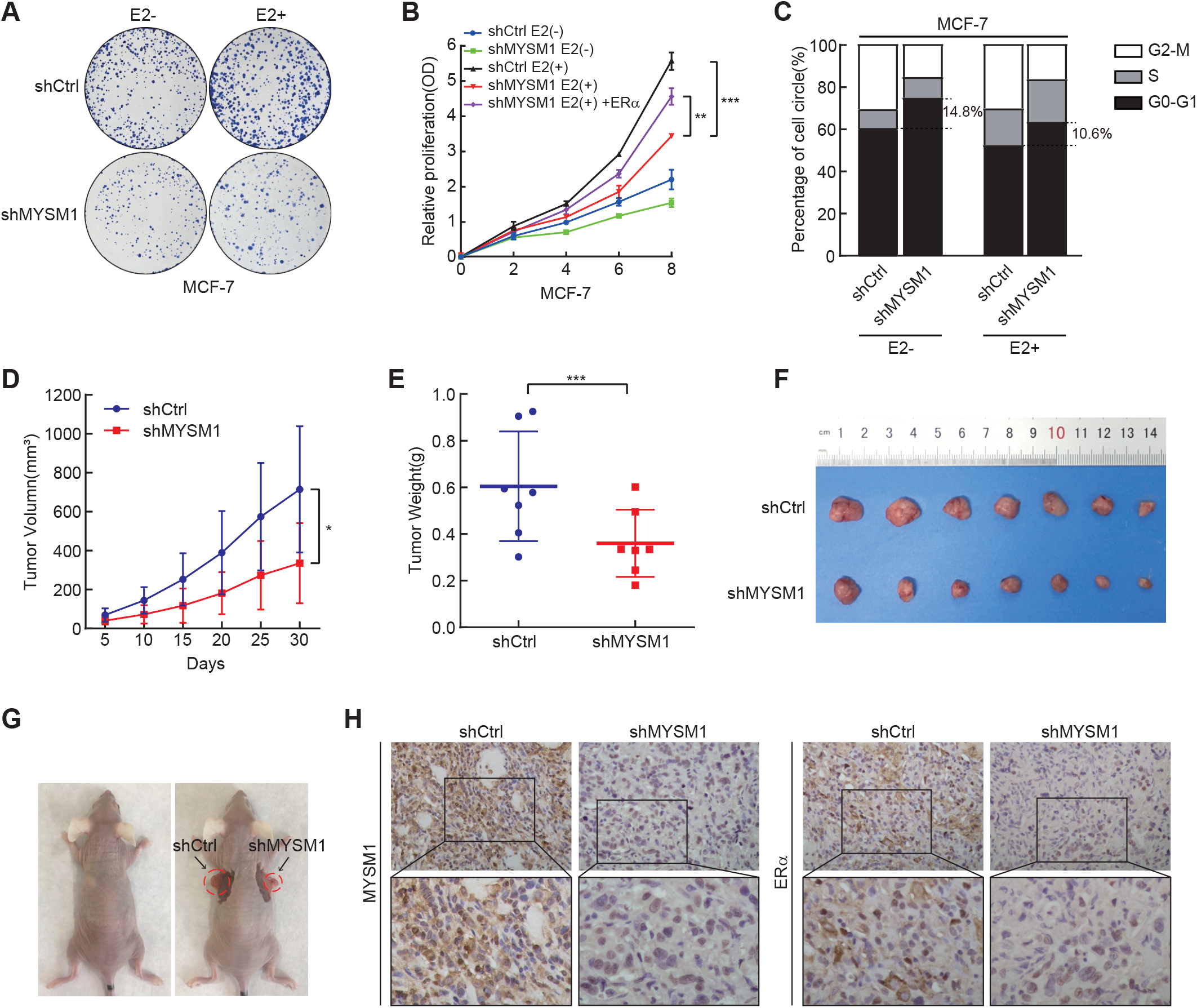
Disrupting MYSM1 dampens breast cancer cell growth. A. Colony formation assay of shCtrl and shMYSM1 stably expressed MCF-7 cells cultured under vehicle or E2 (100 nM) treatment for 15 days. B. Growth curve showing the effect of MYSM1 knockdown on MCF-7 cell proliferation with or without E2 (100 nM). Total cell viability was assessed every other day by MTS assay. \*\**P* < 0.01, \*\*\**P* < 0.001 (mean ± SD; Student *t*-test). C. Representative histogram displaying percentage of the cell population in G0-G1, S, and G2-M phases in MCF-7 cells carrying shCtrl or shMYSM1 under E2 stimulation or not, as detected by flow cytometry. D. Tumor xenografts were generated by injecting MCF-7 cells infected with shCtrl and shMYSM1 subcutaneously in female *BALB/c* mice. The average tumor volume was measured at the indicated time point (mean ± SD; Student *t*-test; n=7). E. Tumor weights of the shCtrl and shMYSM1-group were measured 30 days after cell injection. F, G. Representative images of the dissected tumors harboring shCtrl and shMYSM1-expressed MCF-7 cells. H. Paraffin sections of the nude mice tumors were conducted to immunohistochemistry (IHC) staining with anti-MYSM1 and anti-ERα. The representative pictures were taken at 20× magnification of microscopic field.

To test the functional of MYSM1 on BCa cell growth *in vivo*, MCF-7 cells infected with shMYSM1 or negative control lentivirus (shCtrl) were individually implanted subcutaneously into flank sides of 4-week-old female *BALB/c* nude mice (n = 7) armpit. We monitored tumor growth and measured tumor sizes every 5 days after injection. Compared with shCtrl-MCF-7 cells, shMYSM1-MCF-7 cells carried decreased tumor growth rates (Fig 5D). In accordance with the growth curve, tumors originating from shMYSM1-MCF-7 cells exhibited lower weights and smaller sizes than those from the control (Fig 5E-G). Moreover, immunohistochemical (IHC) analysis was performed to examine the expression in ERα and its target expression. The results revealed that loss of MYSM1 dramatically decreased ERα and c-Myc protein expression in xenograft tumor tissue (Fig 5H and Fig EV4F). Taken together, these results demonstrated that MYSM1 depletion suppressed the breast cancer cell growth in mice.

### MYSM1 depletion facilitates the sensitivity of ERα-positive breast cancer cells to antiestrogen treatment

Estrogen inhibitors or ERα antagonists dependent on the estrogen-ERα axis have ruled supremely for decades to treat ERα-positive breast cancer. Letrozole belonging to aromatase inhibitors (AIs), Tamoxifen acting as a selective ERα modulator (SERM), and Fulvestrant as a selective ERα degrader (SERD) are used as the putative ERα antagonists. Having established that MYSM1 participates in modulation of ERα signaling pathway, we postulate that MYSM1 might be implicated in endocrine resistance process. We thus turned to collect the clinical ERα-positive breast cancer patients with AI adjuvant treatment. Western blot was performed to detect MYSM1 expression in biopsy samples of these patients before and after AI adjuvant treatment. According to Ki67 index and clinical imaging features, these patients were separated into responders and non-responders. The grayscale analysis in two cohorts as indicated showed that MYSM1 protein was up-regulated in non-responders versus responders (Fig 6A).

**Figure 6.**
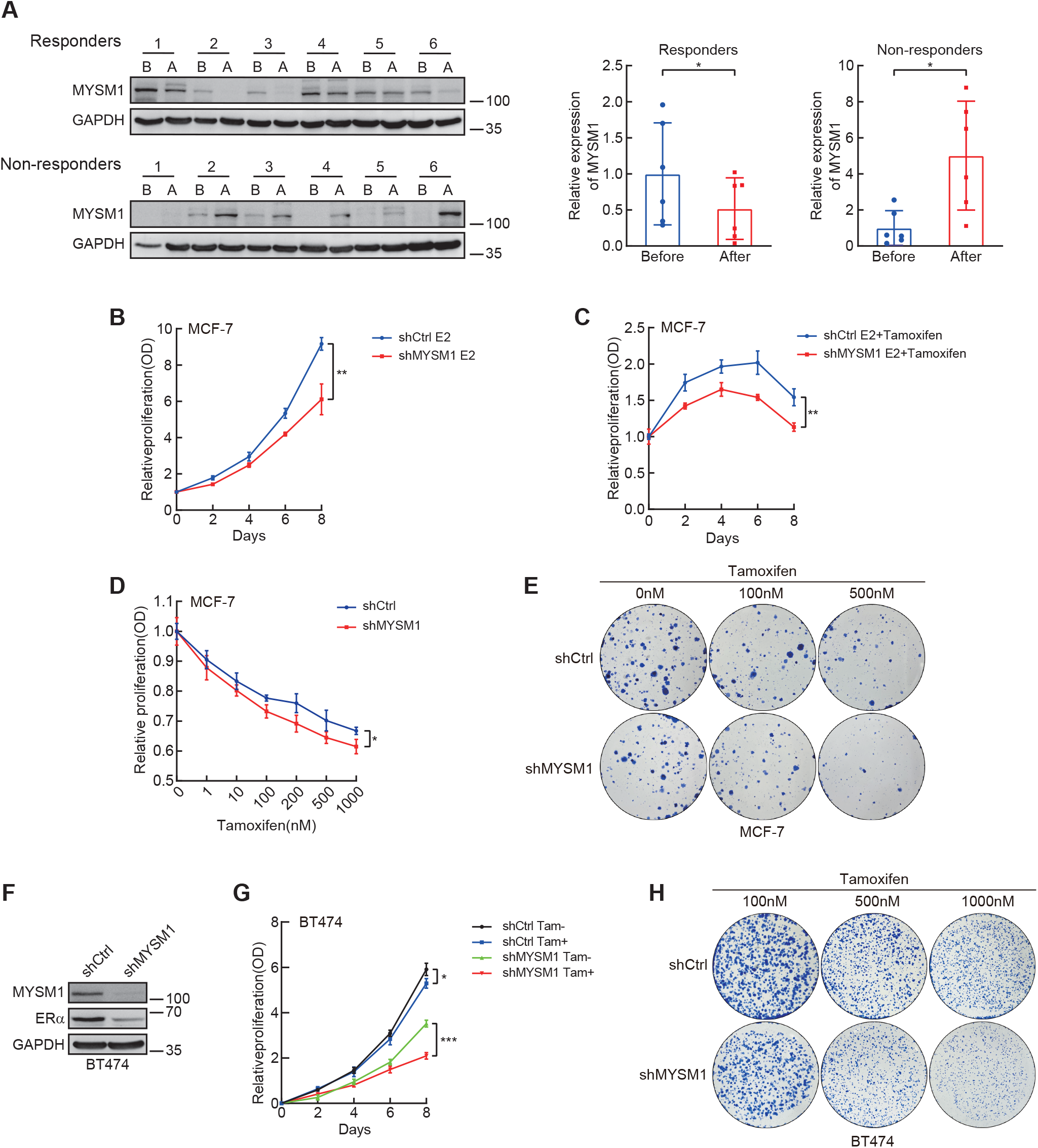
MYSM1 depletion subjects ER-positive breast cancer cells to endocrine treatment. A. ER-positive BCa patients that received AI adjuvant treatment were divided into 2 groups named “Responders” and “Non-responders”. Clinical biopsy samples before and after treatment were examined by western blot. Semiquantitative analyses are presented as relative expression of MYSM1 normalized to that of the before-group. \**P* < 0.05 (mean ± SD; Student *t*-test; n=6). B, C. Growth curve showing the effect of MYSM1 knockdown on MCF-7 cell proliferation with Tamoxifen (500nM) absence (B) or presence (C). Total cell viability was assessed every other day by MTS assay. \*\**P* < 0.01 (mean ± SD; Student *t*-test). D. A cellular viability detection in MYSM1-deletion MCF-7 cells that incubated in various concentrations of Tamoxifen for 7 days. E, H. The panels show colony-formation assay conducted in MCF-7 or BT474 cells infected with lentivirus expressing shCtrl/shMYSM1. Cells in each panel were treated with different doses of Tamoxifen for 15 days before fixation and R250 staining. F. Western blot experiment to validate MYSM1-knockdown efficiency in BT474 cells. G. The line chart renders the relative proliferation of shMYSM1-BT474 cells compared to shCtrl-BT474 cells in the stimulation of Tamoxifen or not. \**P* < 0.05, \*\*\**P* < 0.001 (mean ± SD; Student *t*-test).

To examine the effect of MYSM1 expression on the antiestrogen effects, cell viability was evaluated in MYSM1-silencing MCF-7 cells with ERα antagonist treatment. Compared with the group of E2 existence only, MYSM1 depletion exacerbates cell death with an extra addition of appropriate concentration of Tamoxifen, Fulvestrant, or Letrozole. These results suggest that MYSM1 may confer the resistance to antiestrogen treatment (Fig 6B and C, and Fig EV5A and B). Besides, cell growth curves and colony formation proved that MYSM1 depletion made cancer cells more susceptible to the drug toxicity effects under the condition of higher dosage of specific endocrine drugs (Fig 6D and E, and Fig EV5C-F). Furthermore, BT474 cells owing intrinsic Tamoxifen resistance were also used to construct cell lines stably infected with shMYSM1 (Fig 6F). Fig 6 G and H showed that MYSM1 deficiency facilitated the sensitivity to Tamoxifen treatment in BT474 cells.

### Expression of MYSM1 is upregulated in clinical Breast Cancer tissues

Having shown that MYSM1 acting as the novel ERα co-activator is involved in the maintenance of ERα stability via its deubiquitinase activity. We thus wanted to estimate the correlation between MYSM1 and ERα in clinical BCa samples. Western blot was performed with specimens from 30 pairs of fresh ERα-positive breast cancer tissues and the adjacent benign mammary tissues. MYSM1 protein is prominently elevated in breast cancer cohort (Fig 7A and B). Moreover, its expression was positively correlated with that of ERα (Fig 7A and C). To sketch out the clinical relevance of MYSM1 in breast cancer, we took advantage of the commercial tissue microarrays as well as some paraffin sections owing detailed patient information sponsored by hospital. The results from IHC staining with the anti-MYSM1 antibody demonstrated that the expression of MYSM1 gradually increased as the BCa proceeded into higher grade (Fig 7D and E). The statistical analysis of clinicopathological characteristics proposed a tight bond in terms of MYSM1 expression with histological grade, ERα status, and HER2 status (Table 1). Finally, patients were split into two groups based on the median MYSM1 expression value to calculate the survival rates. The data showed that higher expression of MYSM1 exhibited the poor overall survival, implying that MYSM1 may act as an indicator in breast cancer prognosis (Fig 7F).

**Table 1.**
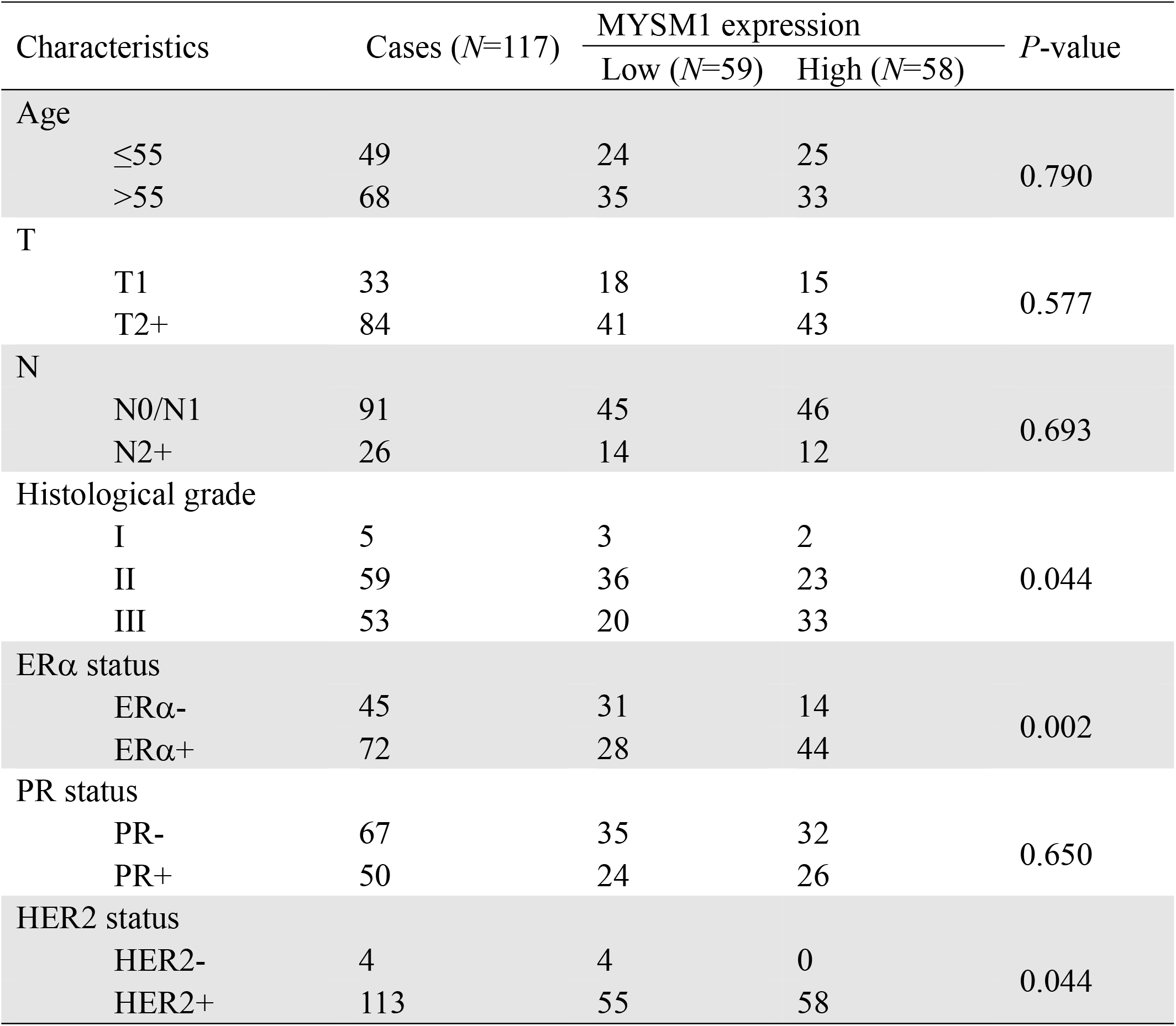
Relationship between MYSM1 expression and clinical pathologic features in breast cancer.

| Characteristics | Cases (N=117) | MYSM1 expression |  | P-value |
| --- | --- | --- | --- | --- |
|  |  | Low (N=59) | High (N=58) |  |
| Age |  |  |  |  |
| ≤55 | 49 | 24 | 25 | 0.790 |
| >55 | 68 | 35 | 33 |  |
| T |  |  |  |  |
| T1 | 33 | 18 | 15 | 0.577 |
| T2+ | 84 | 41 | 43 |  |
| N |  |  |  |  |
| N0/N1 | 91 | 45 | 46 | 0.693 |
| N2+ | 26 | 14 | 12 |  |
| Histological grade |  |  |  |  |
| I | 5 | 3 | 2 | 0.044 |
| II | 59 | 36 | 23 |  |
| III | 53 | 20 | 33 |  |
| ERα status |  |  |  |  |
| ERα- | 45 | 31 | 14 | 0.002 |
| ERα+ | 72 | 28 | 44 |  |
| PR status |  |  |  |  |
| PR- | 67 | 35 | 32 | 0.650 |
| PR+ | 50 | 24 | 26 |  |
| HER2 status |  |  |  |  |
| HER2- | 4 | 4 | 0 | 0.044 |
| HER2+ | 113 | 55 | 58 |  |

**Figure 7.**
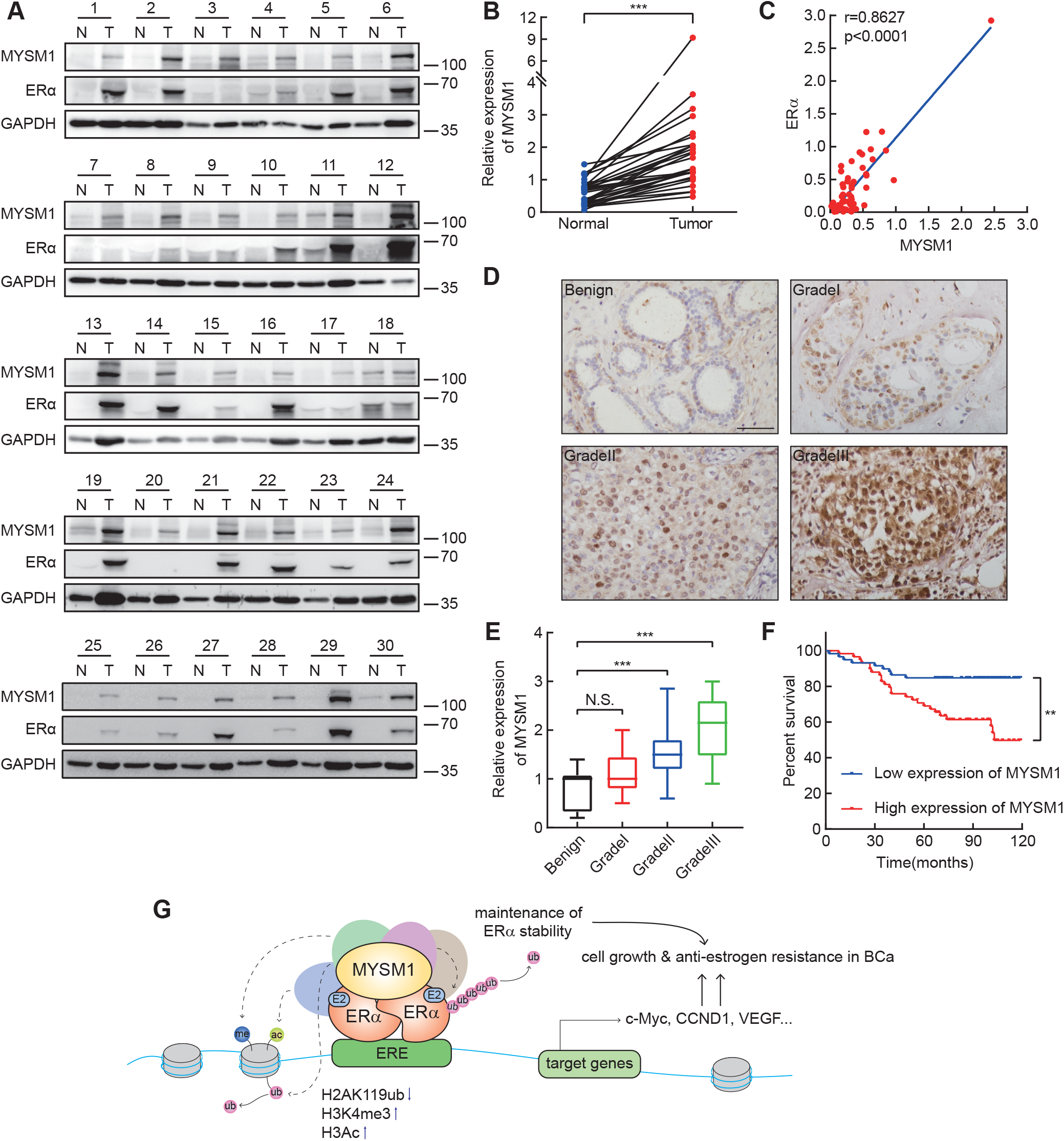
Expression of MYSM1 is upregulated in clinical Breast Cancer tissues. A. The protein expression of MYSM1 and ERα in 30 presentative pairs of primary BCa (T) and adjacent non-cancerous tissues (N). B. Grayscale analysis of MYSM1 level was conducted using GAPDH as the internal control and the differential expression of MYSM1 in T versus N was delineated. \*\*\**P* < 0.001, student *t*-test. C. The level of MYSM1 is positively correlated with ERα. Semiquantitative expression of MYSM1 and ERα from the 30-pair samples were statistically analyzed. The relative level of MYSM1 was plotted against that of ERα. D. Representative MYSM1 IHC images of primary ER-positive breast tumors in different grades. Scale bars, 100 µm. E. Statistical quantifications of MYSM1 expression in BCa samples as differentiation progresses. \*\*\**P* < 0.001 and N.S. stood for no significant (mean ± SD; Student *t*-test; n=149). F. Higher MYSM1 level predicts a poor clinical outcome in BCa cases. Median expression score of MYSM1 was used as cutoff to evaluate the overall survival by Kaplan-Meier (KM) method. \*\**P* < 0.01 G. Model of MYSM1 regulating the ERα signaling in ER-positive breast cancer.

## Discussion

The ERα signaling acts as an obligate component in female mammary growth and development. It requires strict supervision to prevent against blunted mammary glands morphogenesis or the tumorigenic processes (Rusidze et al, 2021). The extraordinarily higher expression of ERα protein in breast cancer tissues reflects a specific genomic environment and intricate signaling networks in tumor cells. In this study, we identified a histone deubiquitinase, MYSM1 as a novel ERα co-activator in breast cancer. Our results have demonstrated that MYSM1 is recruited together with ERα to the promoter region of ERα target genes, leading to epigenetic modulation of H2Aub, H3K4me3, and H3Ac levels. On the other hand, we provided the evidence that MYSM1 is involved in maintenance of ERα stability via reduction of Lysine 48 (K48) and K63-linked poly-ubiquitination on ERα. Thus, MYSM1 enhances ERα action via histone and non-histone deubiquitination to promote cell proliferation and antiestrogen insensitivity in breast cancer progression. Our study suggests that MYSM1 acts as an efficiency co-activator of ERα, exerting epigenetic modifier and ERα stability maintainer to enhance antiestrogen insensitivity in ERα positive breast cancer (Fig 7G).

Modification of histone tails by epigenetic regulators results in reprogramming of chromatin landscape to regulate gene transcription (Zhao et al, 2021). Histone H2A ubiquitination is a highly conserved histone modification that compacts chromatin structure and decreases chromatin accessibility, thereby reducing the specific affinity of transcription factors to inhibit transcriptional regulation of non-lineage specific target genes(Barbour et al, 2020; Higashi et al, 2010; Tamburri et al, 2020). Uncontrolled H2Aub deposition leads to genomic instability and is prevalent in cancer development(Tamburri et al, 2021). MYSM1 as a histone H2A deubiquitinase has been reported to activate the transcription of its downstream genes, including CDH1, c-MET, or genes encoding ribosomal proteins via histone H2A deubiquitination in colorectal cancer, melanoma, and B cell lymphoma (Chen et al, 2021; Lin et al, 2021; Wilms et al, 2017). Consistent with these studies, our results indicate that MYSM1 enhances ERα-induced transactivation by eliminating H2Aub deposition. In addition, MYSM1 appears to motivate the accumulation of active epigenetic marks H3K4me3 and H3Ac. In fact, previous studies have demonstrated that MYSM1 interplays with p/CAF to form a co-regulatory protein complex that stepwise synergizes histone ubiquitination and acetylation for AR-dependent gene activation in prostate cancer cells(Zhu et al, 2007). MYSM1 may be a component of protein complexes that regulate ERα target genes cross-talking with other histone modifiers in a serial and combinatorial manner in the breast cancer cellular context, which is coincident with our predicted MYSM1-relevant PPI networks from the STRING database (Fig EV1C).

By cleaving ubiquitin chains from non-histone substrates, deubiquitinase could accelerate tumor development by modulating the expression, activity, and localization of many substrates (Pal et al, 2014). ERα has been shown to be a substrate of deubiquitinase (Pesiri et al, 2016). A few DUBs exert their pro-oncogenic function by triggering ERα deubiquitination to maintain the stability of ERα in BCa (Cao et al, 2021; Tang et al, 2021; Wang et al, 2020; Xia et al, 2021; Xia et al, 2019). Furthermore, DUB inhibitors targeting USP14 and UCHL5 downregulate ERα expression and induce cancer cell apoptosis, providing a prospective strategy for ER-positive BCa treatment (Xia et al, 2018). Due to the characteristic of a higher turnover rate and greater homeostasis demand of ERα protein in breast cancer cells, ERα-targeting DUBs have become critical targets for novel cancer therapeutics discovery (Xiao et al, 2016). As a ubiquitin-specific protease, MYSM1 was reported to remove K63-linked polyubiquitin chains of TRAF3, TRAF6, RIP2, and STING, resulting in the collapse of the complex scaffolds and attenuated complex assembly, thereby attenuating the associated signaling pathways (Panda & Gekara, 2018; Panda et al, 2015; Tian et al, 2020b). However, its catalytic activity towards polyubiquitin chains of different geometric configurations on different substrates remains unclarified. Herein, our data demonstrated that MYSM1 is involved in ERα deubiquitination and maintains ERα stability. We further provided the evidence to show that MPN domain of MYSM1 is required for maintenance of ERα stability, suggesting MYSM1 inhibits ERα degradation via its catalytic activity. Further study indicated that MYSM1 removes the K48- and K63-linked polyubiquitination on ERα. K48-linked polyubiquitination induces degradation of proteasomal substrates, and K63-linked polyubiquitination mainly modulates non-proteolytic cellular processes, such as signal transduction. One study found that K63 chains conjugated to TXNIP could attract K48 ligases to trigger K48/K63 branched chains formation and regulates proteasomal degradation, providing a new insight for heterogenous ubiquitin chains to convert a non-degradative ubiquitin into a degradation signal (Ohtake et al, 2018). So far, it’s unclear whether branched chains could be assembled on ERα protein or the exact function of K63-linked polyubiquitination on signal transduction of ERα pathway. The branched chains represent an increasing level of attached ubiquitin concentrating closely to the substrate for a higher affinity of effectors towards ubiquitin. That could explain the extremely powerful proteolytic signal of these conjugates. The multiple blocks at both types of K48- and K63-linkages by MYSM1 would makes its pro-stabilizing effect on ERα more intense based on the assumption of ERα degradation through K48/K63 branched chains.

Aberrant activation of ERα signaling is a common incentive of endocrine resistance in BCa, which may be induced by mutations in *ESR1* gene, peculiar post-translational modification of ERα protein, abnormal expression of ERα co-regulators, or compensatory cross-talk between ERα and parallel oncogenic signaling pathways (Belachew & Sewasew, 2021; Hanker et al, 2020). These aspects result in the unconventional expression of ERα signature genes involved in diverse biological processes to oppose endocrine therapy (Louie et al, 2010; Miller et al, 2011). Herein, we provided the evidence to show that MYSM1 co-activates ERα action via histone and non-histone manner to confer antiestrogen insensitivity in breast cancer. MYSM1 up-regulated ERα-induced transcriptional activity of target genes, such as c-Myc, VEGF, and CCND1, which play crucial roles in endocrine resistance (Fig 2 and 4). MYSM1 was higher expressed in non-responders after AI adjuvant treatment in ERα positive BCa samples (Fig 6A). The influence of MYSM1 on the effect of antiestrogen treatment demonstrated that MYSM1 depletion increased the sensitivity of breast cancer-derived cells to the treatment of antiestrogen drugs (Fig 6). On the other hand, we also observed that MYSM1 accelerates cell proliferation even in the absence of E2, suggesting that MYSM1 may also participate in regulating non canonic ERα signaling pathway independent on E2 treatment (Fig 5A and B). Thus, in this study, we identified MYSM1 as a novel ERα co-activator is involved in histone H2A and ERα deubiquitination, thereby up-regulating ERα-mediated transactivation and ERα itself stability maintenance to confer antiestrogen resistance in ERα positive BCa.

Collectively, our data have demonstrated the function of MYSM1 on maintenance of ERα stability and modulation of ERα action to confer antiestrogen insensitivity in ERα-positive breast cancer, providing a potential therapeutic target for endocrine resistance in ERα positive BCa.

## Materials and methods

### Drosophila strains and genetics

Fly stocks were maintained on cornmeal sucrose-based media at 25°C. The human ERα and MYSM1 cDNA cloning products were introduced in a modified pCaSpeR3 vector with upstream active sequence (UAS) promoter to generate the UAS-linked ERα/MYSM1 expression construct. For the reporter construct, ERE sequence copies were integrated upstream of the GFP reporter genes controlled by the TATA promoter box. The UAS-ERα/MYSM1 expression and ERE-GFP reporter constructs were sent to EMBL *Drosophila* Injection Service for transgenic flies generation. The CG4751 deletion mutant (CG4751^-/+^) were obtained from Bloomington *Drosophila* Stock Center. To examine the effect of CG4751 loss or MYSM1 gain of function on ERα-dependent transactivation of reporter gene, we crossed the male F0 owing hemizygous mutants (deletion or overexpression) with Gal4-UAS ERE-GFP female flies. F1 progeny possessing the mutant allele and mosaic red eye were carried out for eye disc histology analysis as previously described(Sun et al, 2016).

### Plasmids and antibodies

The antibodies used in this study were: anti-MYSM1 (Abcam, cat# ab193081), anti-ERα (Cell Signaling, cat# 8644), anti-c-Myc (Proteintech, cat# 10828), anti-VEGF (Proteintech, cat# 19003-1-AP), anti-Cyclin D1 (Cell Signaling, cat# 2978), anti-GAPDH (ABclonal, cat# AC036), anti-FLAG, anti-His and anti-HA (GNI), anti-H2AK119Ub (Cell Signaling, cat# D27C4), anti-H3K4me3 (Sigma-Aldrich, cat# 07-473), anti-H3Ac (Sigma-Aldrich, cat# 06-599).

Full-length cDNA for MYSM1 was amplified by PCR using the cDNA library as a template. The coding sequences of various MYSM1 mutants amplified using the primers listed in Supplementary Table 1. The PCR products for MYSM1 wild-type and a series of mutants were cloned into a p3×FLAG-CMV-10 expression vector to generate MYSM1-FL, MYSM1-ΔMPN, MYSM1-ΔSANT and MYSM1-ΔSWIRM. Other plasmids were identical to our previous studies.

### Cell lines and culture conditions

All cell lines were obtained from the ATCC (American Type Culture Collection). MCF-7 and HEK293 were cultured in DMEM medium, supplemented with 10% fetal bovine serum (FBS; Gibco). T47D and BT474 were cultured in or RPMI-1640 medium, supplemented with 10% fetal bovine serum (FBS; Gibco). For estrogen-starving conditions, cells were grown in phenol red-free DMEM or RMPI-1640 containing 5% charcoal-treated serum. All cells were maintained in a humidified incubator at 37 °C and 5% CO2.

### siRNA and lentivirus

siRNA control (siCtrl) and siRNA against the gene encoding MYSM1 were purchased from Sigma Aldrich. The RNAi nucleotides were transiently transfected in cells using jet-PRIME transfection reagent (Polyplus) following the manufacturer’s instructions. Sequence of siMYSM1: 5’-CAAAUGCGGUCUGGAUAAAdTdT-3’. Sequence of siCtrl: 5’-UUCUCCGAACGUGUCACGUdTdT -3’. For lentivirus-delivered RNAi, negative control (shCtrl) and shRNA against MYSM1 (shMYSM1) lentivirus targeting the same sequence as siMYSM1 as above were purchased from Shanghai GeneChem Company. Two days after lentivirus infection, puromycin was added into the medium at a concentration of 3 µg/ml to select stably transduced cells.

### Western blot and Co-Immunoprecipitation (Co-IP)

Western blot was performed by the standard process as previously described(Zhang et al, 2022). Briefly, samples were lysed in lysis buffer containing protease inhibitor and the supernatants harvested from centrifugation were resolved by SDS-PAGE. After electrophoresis, proteins were transferred to PVDF membrane, probed with primary antibodies overnight at 4 °C, and incubated with the appropriate secondary antibody for 1 h at room temperature. The respective protein bands were visualized by ECL solution. For immunoprecipitation, experiments were performed on the basis of the manufacturer’s instructions. Supernatant cell lysates were incubated with Protein A/G agarose gel and primary antibodies under constant rotation overnight, then the mixture was rinsed three times before analyzed by immunoblotting.

### Ubiquitination assay

The plasmids were transiently transfected into MCF7 or HEK293 cells for 48 h, incubated with 10 μM MG132 for 6 h. The cells were collected and lysed in the denature lysis buffer. The His/HA-ubiquitinated ERα protein was purified and immunoblotted with anti-His or anti-HAantibodies. HA-tagged different ubiquitin mutants, including K0 (lysine less), K48 (only K48-linked-Ub), and K63 (only K63-linked-Ub), were used in this study as indicated.

### GST-pull down

*E. coli* cells expressing GST-conjugated ERα-AF1 and ERα-AF2 protein were lysed to get the purified protein. Glutathione-Sepharose beads (GE Healthcare) coupled with either GST or with the GST fusion protein (GST ERα-AF1 and GST ERα-AF2) were incubated with the *in vitro* transcripted and translated FLAG-MYSM1. After rinsing the beads three times, the proteins bound to the beads were detected by western blot and stained using using Coomassie Brilliant Blue R-250.

### Immunofluorescence (IF)

Cells were fixed at room temperature with 4% paraformaldehyde for 20 min, permeabilized in PBS containing 0.05% Triton X-100 for 20 min, blocked with 1% donkey serum albumin, and incubated with primary antibodies (1:100 dilution) in a humid chamber overnight at 4°C. After three times washing, specimens were incubated with Alexa Fluor conjugated secondary antibody (Jackson ImmunoResearch Laboratories Inc) for 1 h followed by DAPI (Roche) staining for 30 min at room temperature and then subjected to confocal microscopy.

### Luciferase dual-reporter assays

4 h after transiently transfection of wild-type or mutant MYSM1 expression constructs, along with ERα, ERE-tk-Luc, and an internal control plasmid of Renilla luciferase (pRL), cells were cultured in medium containing 5% charcoal-treated serum in the presence of E2 (100 nM). 24 h later, cells were lysed for luciferase activity detection by a Promega dual-luciferase reporter assay system.

### RNA isolation and quantitative real-time PCR (qPCR)

MCF-7 and T47D cells were harvested and total RNA was extracted with RNA Trizol (TAKARA) following the manufacturer’s recommendations. 1 μg of total RNA was reversely transcribed into cDNA using the PimeScript RT-PCR kit (TAKARA) and analyzed by qPCR on LightCycler96 system (Roche) with the SYBR premeraseTaq kit (TAKARA). Amplified mRNA levels were normalized to 18s mRNA. Primers used were listed in Supplementary Table 2.

### Chromatin immunoprecipitation (ChIP) and ChIP re-IP

Cells transfected with siCtrl or siMYSM1 were cultured in phenol red-free medium with 5% charcoal-treated serum for 2 days followed by 6 h E2 (100 nM) or equal EtOH stimulation. Cells were cross-linked with 1% formaldehyde at room temperature before glycine (0.25 M) quenching. After cell collection, lysis buffer was added and the mixture was sonicated on ice. Sheared-chromatin solutions were separated by centrifugation at 12000 rpm 15 min at 4 °C and an eighth of the supernatant was kept for input. Concurrently, the remaining supernatant was immunoprecipitated using specific antibodies overnight. After that, specimens were incubated with protein A-sepharose beads for 4 h on a mixing platform before sequentially beads-washing by low salt buffer, high salt buffer, LiCl buffer, and TE buffer. The protein-DNA complexes were eluted by elution buffer followed by crosslink reversal and proteinase K digestion. Finally, the purified DNA was precipitated in absolute alcohol and resuspended in TE buffer. qPCR was performed to examine the precipitated genomic DNA samples(Nelson et al, 2006). The sequences of the primers described in Supplementary Table 3. For ChIP re-IP, the cross-linked immunocomplex was eluted from the first ChIP by 10 mM dithiothreitol incubation at 37 °C for 30 min and diluted 50-fold in re-ChIP buffer. The products were subjected to the ChIP procedure again with the indicated antibodies.

### Cell viability, colony formation and flow cytometric analysis

Cells infected with shCtrl or shMYSM1 lentivirus were seeded in 96-well plates at 3000 cells per well, in triplicate, complemented with estradiol, Tamoxifen (Abmole, Cat# M7353), Fulvestrant (Abmole, Cat# M1966) or Letrozole (Abmole, Cat# M3699) for 7 days. Viable cells at different time points were measured by MTS assay (Promega, Cat# G3580) with the absorbance at a wave-length of 490 nm.

For colony formation assay, cells were planted in 6-well plates and cultured in medium with E2, Tamoxifen, Fulvestrant or Letrozole treatments at stated concentrations for 2 weeks, then were fixed by 4% paraformaldehyde for 15 min before undergoing crystal violet staining of colony number count.

For analysis of cell cycle, cells were harvested with EDTA-free trypsin two days after E2 (100 nM) addition and fixed with 75% ethanol at -20 °C overnight. After ice-cold PBS washing for once the next day, nuclear DNA was stained by propidium iodide for 15 min in a dark room and determined on a flow cytometer (Becton Dickinson).

### Mouse xenograft studies

The animal experiments were approved by Animal Ethics Committee of China Medical University. 5 × 10^6^ MCF-7 cells stably expressed shCtrl and shMYSM1 lentivirus were suspended in 100 μl of culture medium/Matrigel (1:1) (BD Biosciences) and implanted subcutaneously into the bilateral armpits of 4-week-old female *BALB/c* nude mice (Vital River Laboratory). Tumor diameter was measured every 5 days and tumor volume was calculated as follows: *V* = (0.5 × width^2^ × length) mm^3^. Four weeks after inoculation, tumor-bearing mice were sacrificed following the policy for the humane treatment of animals and tumor tissues were isolated, weighing, followed by formalin-fixing and paraffin-embedding for IHC staining.

### Patients and tumor specimens

Human primary BCa tissues and corresponding adjacent tissues were obtained from Liaoning Cancer Hospital and the First Affiliated Hospital of China Medical University. All specimens were collected with patients’ informed consents.

### Immunohistochemical (IHC) analysis

Commercial BCa tissue microarray slides (HBre-Duc150Sur-02) were purchased from Shanghai Outdo Biotech Co., Ltd. (SOBC). Paraffin-embedded tissue sections were deparaffinized and rehydrated before endogenous peroxidase removal by 3% hydrogen peroxide for 15 min. Subsequently, tissues were boiled in citrate buffer (pH 6.0) for antigen retrieval in a pressure cooker followed by protein block with goat serum for 30 min at room temperature. The sections were next incubated with anti-MYSM1 (Sigma-Aldrich, cat# HPA054291, dilution 1:1000), anti-ERα (Cell Signaling, cat# 8644, dilution 1:200), c-Myc (Proteintech, cat# 10828, dilution 1:400) antibodies respectively at 4 °C overnight and with the secondary horseradish peroxidase-labeled polymer anti-rabbit IgG at room temperature for 20 min. Using DAB as a chromogen, the nuclei were counter stained with hematoxylin. Slides were scanned with an Olympus microscope at 20×. The staining scores were evaluated by H score method based on the proportion of positively stained tumor cells (0-100%) and the brown intensity (0-3). The final expression score, which ranged from 0 to 3, was determined by multiplying these two independent indexes.

### Statistics

Statistical analyses were performed using GraphPad Prism 8.0. and the SPSS statistical software program. For all in vitro and animal experiments, two-tailed Student’s *t* test was performed to calculate the *P* value and data in bar graphs represent mean ± SD of at least three technical replicates. Correlation between MYSM1 expression and clinical parameters was determined by chi-square test. Overall survival curves were plotted according to the Kaplan–Meier method with the log-rank test applied for comparison.

## Data availability

Supplementary Data are available at *The EMBO Journal* online. This study includes no data deposited in external repositories.

## Acknowledgments

We appreciate Dr Yunlong Huo for helpful technique assistance. We thank Dr Shigeaki Kato (Soma Central Hospital, Fukushima, Japan) for providing a pERE-tk-Luc reporter vector, expression plasmids for ERα, and its truncated mutants. We thank Dr Yujie Sun (Nanjing Medical University, Nanjing, China) for MCF-7 cells, which were purchased from American Type Culture Collection.

This work was supported by National Natural Science Foundation of China (32170603, 31871286 for Yue Zhao, 81872015, 82273123 for Chunyu Wang,32100440 for Ge Sun); China Postdoctoral Science Foundation (276066) for Ge Sun; Foundation of Liaoning Province of China (LJKZ0756 for Shengli Wang); Local projects supported by the central government (2022JH6/100100035 for Yue Zhao); Foreign expert project of Ministry of Science and Technology (G2022006007L for Yue Zhao).

## Author contributions

RL, GS, BZ, MW, YB, HL, and SW performed the experiments for this work; RL and YZ developed the working hypothesis and scientific concept; RL contributed all *in vitro* and *in vivo* experiments; GS, BZ, and MW contributed *in vitro* experiments and assisted by CW, SW, and KZ; YB, HL, SW performed murine *in vivo* experiments and assisted by RZ, LL, and WL; QZ and YZ contributed mice or reagents. RL and GS designed the experiments. RL and GS analyzed the data. QZ, YZ, and RL wrote and edited the manuscript. LL and WL supervised the project.

## Figure legends

**Figure EV1.**
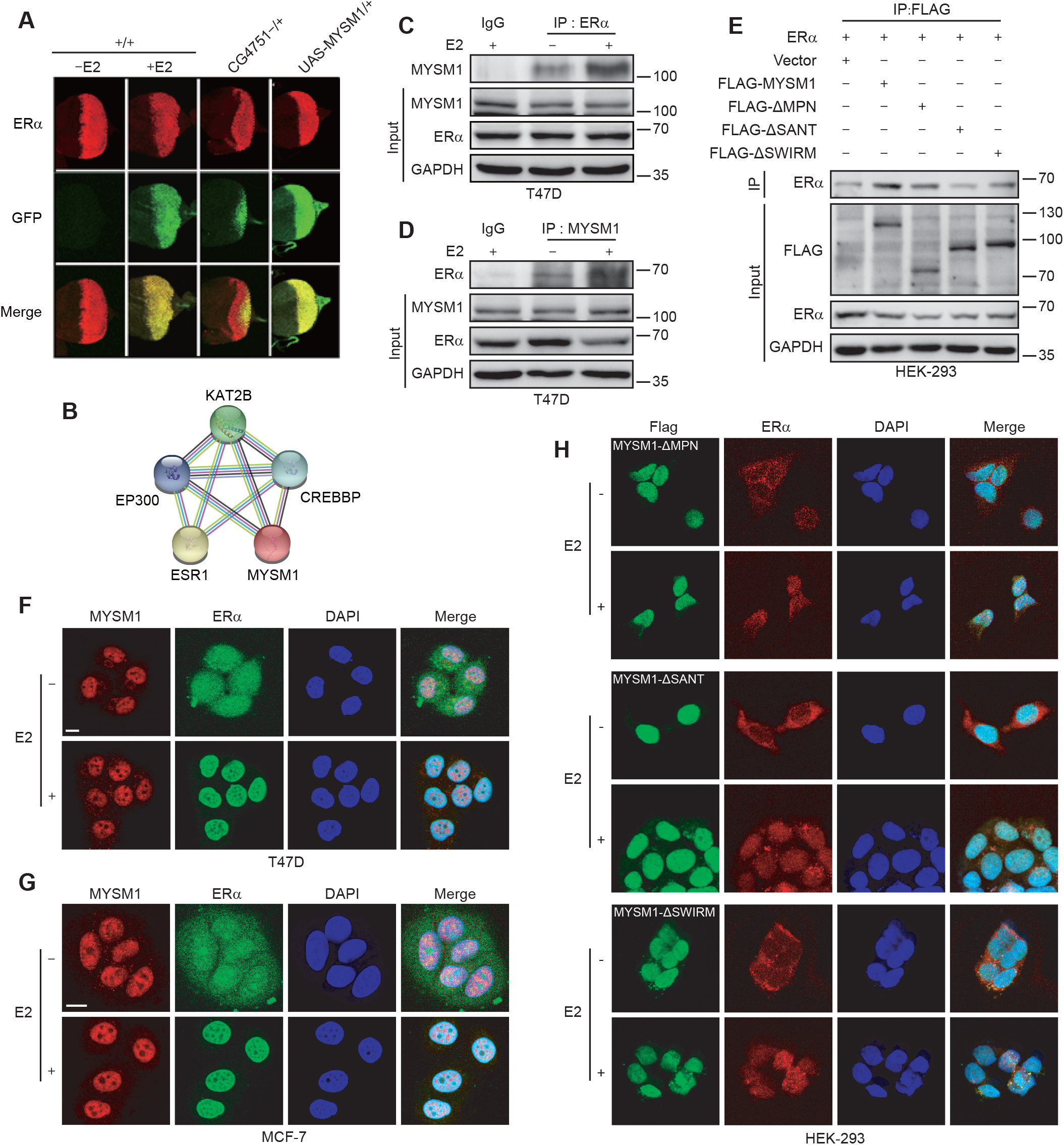
MYSM1 interacts with ERα in breast cancer cells. A. Schematic diagram of the co-regulator screening *Drosophila* model carrying an ERα-mediated gene transcription system. The F0 parental generation with a GAL4-UAS driver targeted ERα expression and an ERE-inserted GFP reporter was crossed with species harboring particular gene X deletion or overexpression mutants. GFP expression changes in the eye discs of F1 progenies were examined to appraise the effects of gene X mutants on ERα-induced transactivation with E2 treatment. B. Evaluation of GFP and ERα level in F1 progeny flies with CG4751 loss of function (lane 3) or MYSM1 gain of function (lane 4) mutants. The lower panels represent merge images. C. The MYSM1-related protein-protein interaction (PPI) networks generated by the STRING online database. D, E. Co-immunoprecipitation conducted in T47D cells to detect the association between endogenous MYSM1 and ERα in response to E2 treatment. F. HEK293 cells complemented with ERα or the deletion mutants of MYSM1 were lysed. Complexes precipitated by anti-FLAG were purified and immunoblotted with indicated antibodies. G, H. Immunofluorescence showing the co-localization of MYSM1 (red) and ERα (green) in T47D (G) and MCF-7 (H) cells. Scale bars, 10 µm.

**Figure EV2.**
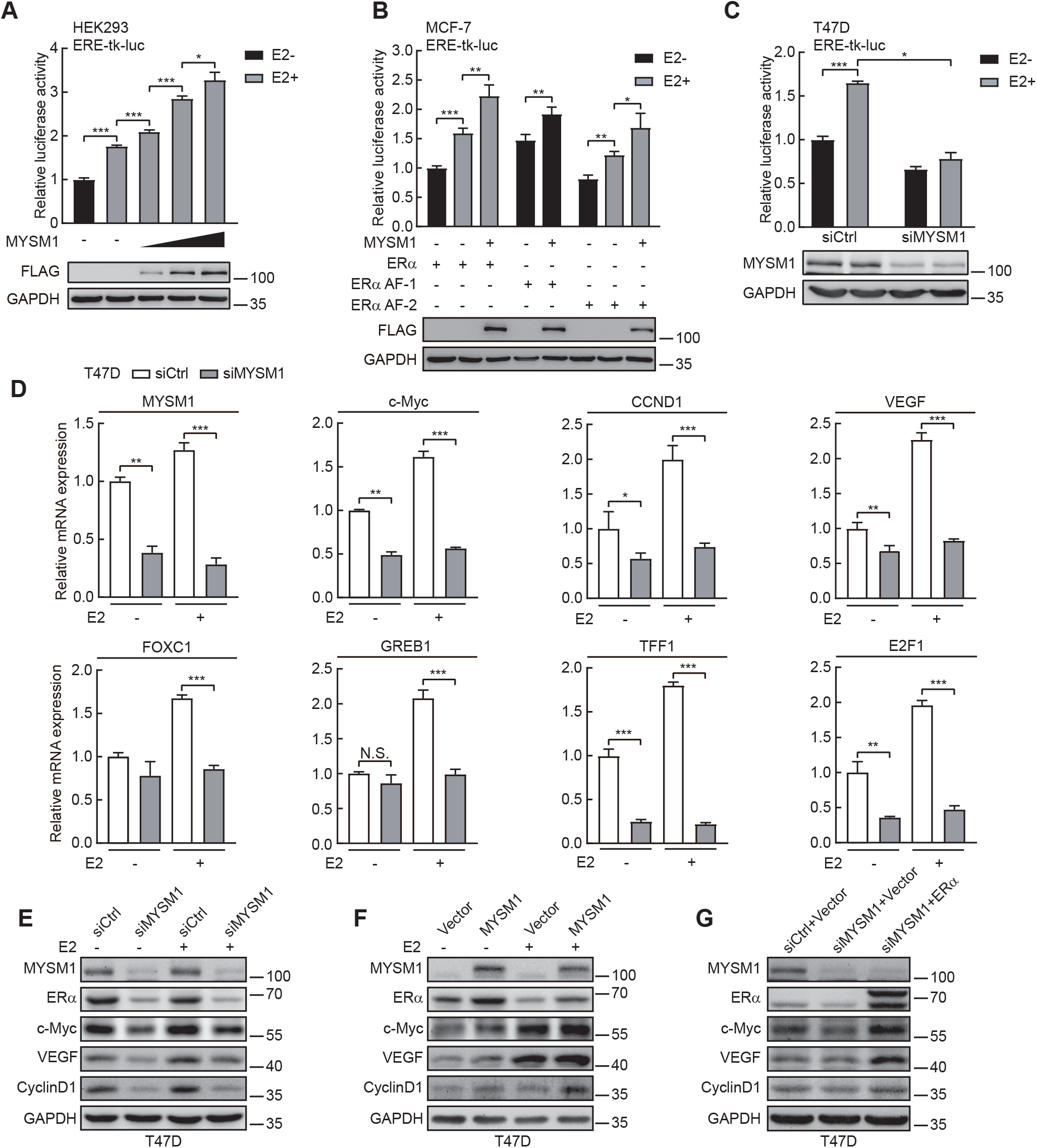
MYSM1 enhances ERα-mediated gene transcription in mammalian cells. A. MYSM1 stimulates ERα-mediated gene transcription in a dose-dependent manner. HEK293 cells were transfected with gradually increased amount of ectopic MYSM1 (0.05μg, 0.1μg, or 0.2μg respectively). MYSM1 expression was examined with anti-B. FLAG by western blot. \**P* < 0.05, \*\*\**P* < 0.001 (mean ± SD; Student *t*-test). B. Relative luciferase activities in MCF-7 cells transfected with ERα full length or truncated mutants harboring ERα AF-1 or ERα AF-2 together with MYSM1 expression plasmid in the presence or absence of E2 (100 nM). The expression of MYSM1 was detected by western blot. \**P* < 0.05; \*\**P* < 0.01; \*\*\**P* < 0.001 (mean ± SD; Student *t*-test). C. Effect of MYSM1 knockdown on ERα-induced transactivation. The relative luciferase values in T47D cells were examined after transient transfection of siCtrl or siMYSM1 followed by ERα expression plasmid. \**P* < 0.05, \*\*\**P* < 0.001 (mean ± SD; Student *t*-test). D. mRNA levels of several ERα target genes in T47D cells with MYSM1-depleted. \**P* < 0.05; \*\**P* < 0.01; \*\*\**P* < 0.001 (mean ± SD; Student *t*-test). E, F. Immunoblot of ERα target gene expression using the indicated antibodies in MYSM1-depleted T47D cells (E) and MYSM1-overexpressed T47D cells (F). G. The loss of ERα and its target genes in MYSM1-depleted T47D cells can be rescued by ectopic ERα expression. T47D cells were transfected with siCtrl or siMYSM1 followed by PcDNA3.1/ERα expression plasmid.

**Figure EV3.**
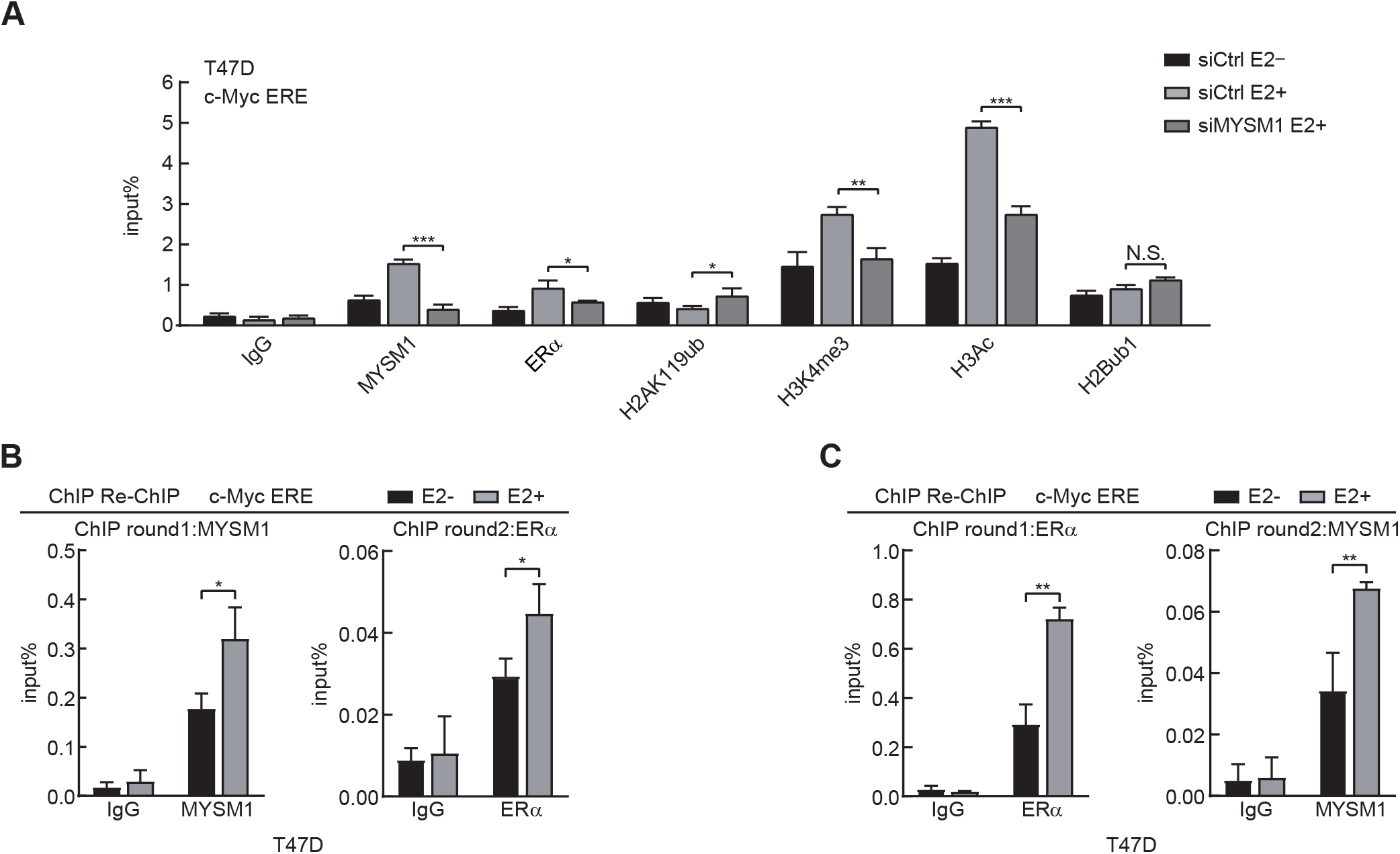
MYSM1 orchestrates ERα occupancy and histone modifications at the locus of ERα target gene. A. ChIP analysis was conducted with specific antibodies in MYSM1-deficiency T47D cells on the promoter region of *c-Myc* with or without E2. \**P* < 0.05; \*\**P* < 0.01; \*\*\**P* < 0.001 (mean ± SD; Student *t*-test). B, C. ChIP re-ChIP assay was performed with the indicated antibodies in T47D cells. \*\**P* < 0.01 (mean ± SD; Student *t*-test).

**Figure EV4.**
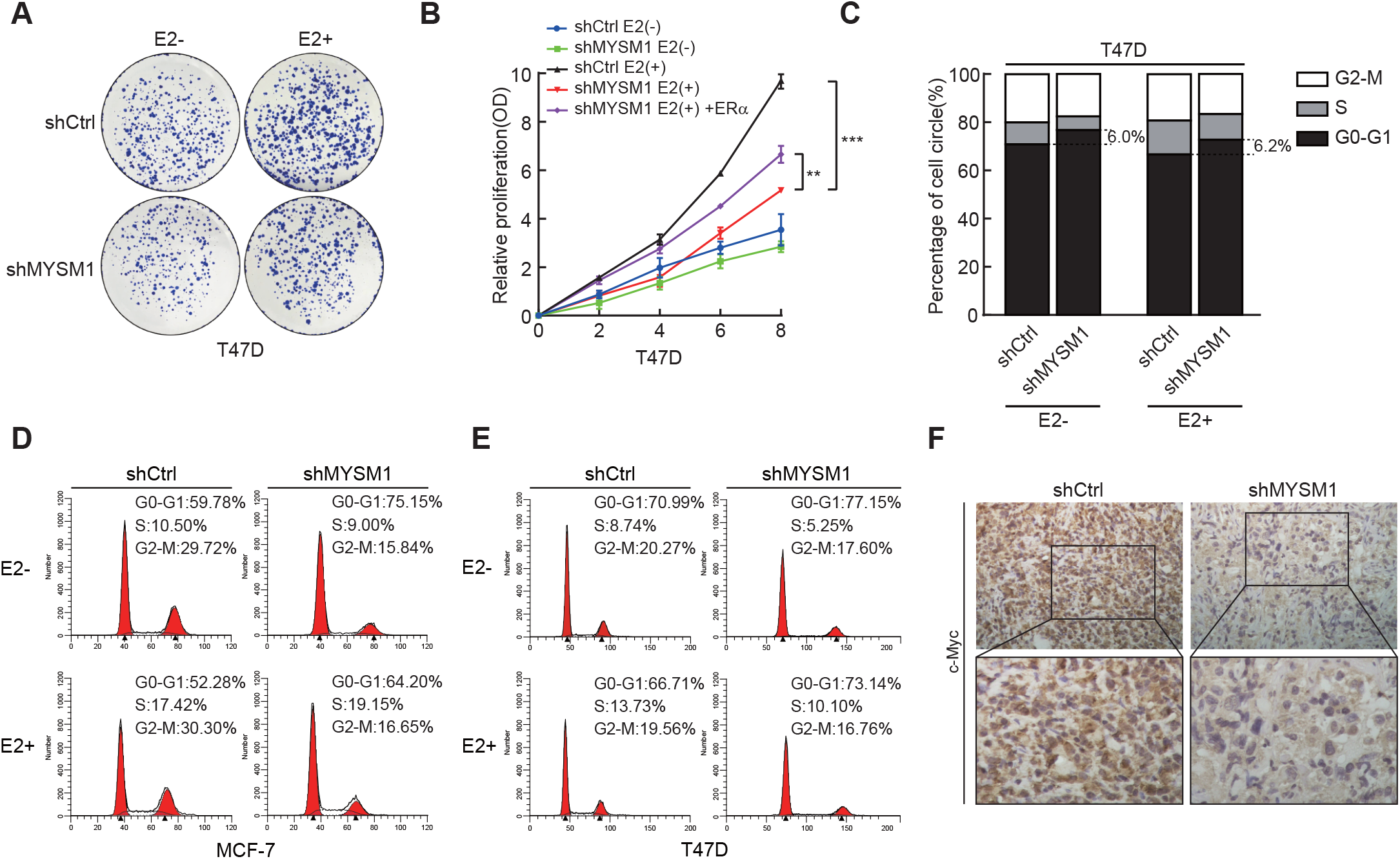
Disrupting MYSM1 dampens breast cancer cell growth. A. Influence of MYSM1 deficiency on T47D cells as illustrated by colony formation. B. Relative proliferation rates of T47D cells carrying shCtrl or shMYSM1 with or without E2 (100 nM) by MTS assay. \*\**P* < 0.01, \*\*\**P* < 0.001 (mean ± SD; Student *t*-test). C-E. Flow cytometric analysis of the cell cycle for MCF-7 (D) and T47D (E) with MYSM1 depletion. The proportion of the T47D cell population in different phases are listed in (C). F. c-Myc expression by IHC analysis in xenograft tumors derived from MCF-7 (shCtrl) group and MCF-7 (shMYSM1) group.

**Figure EV5.**
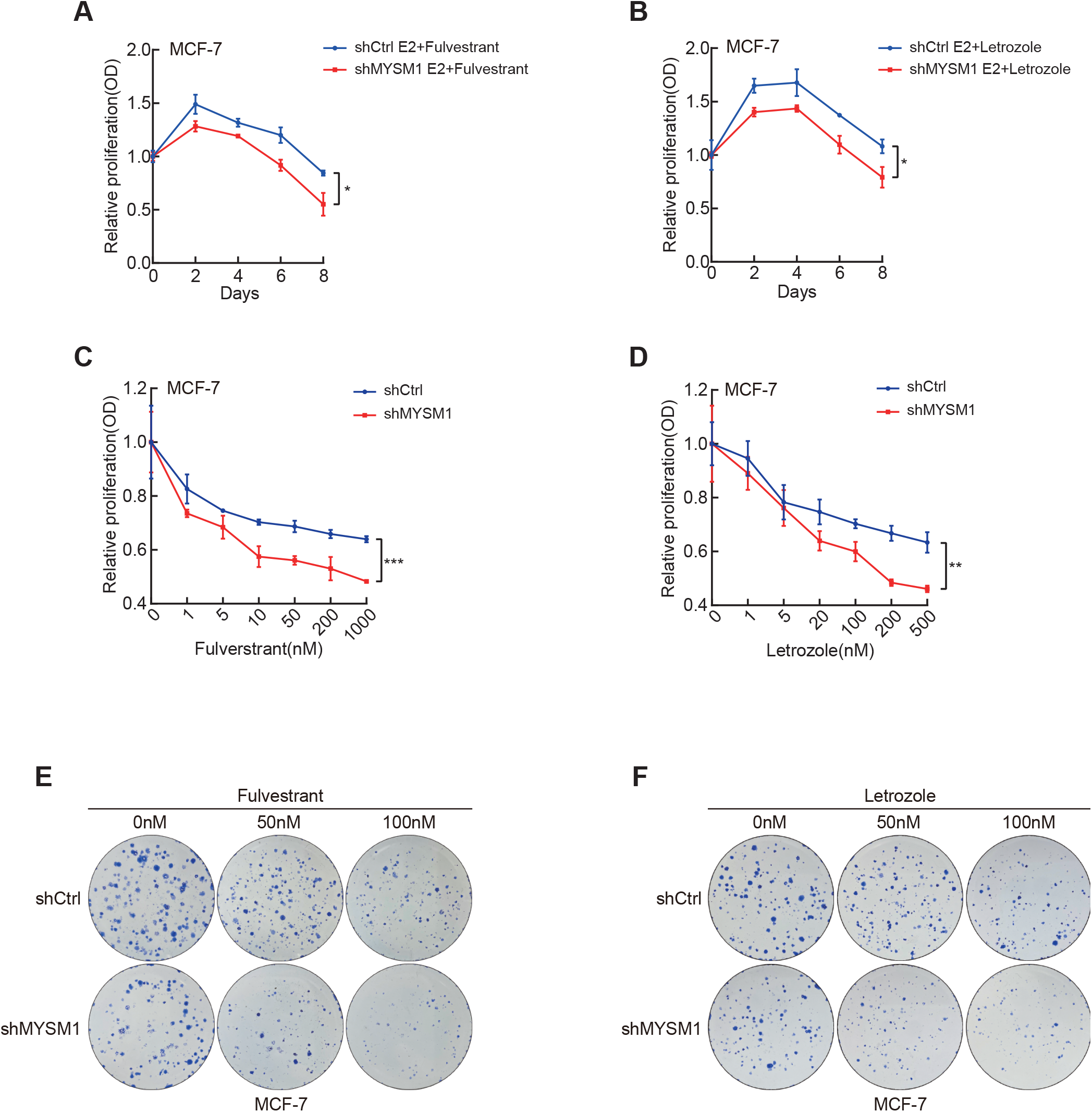
MYSM1 depletion subjects ER-positive breast cancer cells to endocrine treatment. A. MYSM1 protein expression in 6 pairs of fresh ER-positive BCa tissues for the Responders and Non-responders, respectively. B represents cases before AI treatment, A represents cases after AI treatment. B, C. Growth curve showing the effect of MYSM1 knockdown on MCF-7 cell proliferation with Fulverstrant (200nM) (B) or Letrozole (100nM) (C) treatment. Total cell viability was assessed every other day by MTS assay. \**P* < 0.05 (mean ± SD; Student *t*-test). D, E. A cellular viability detection in MYSM1-deletion MCF-7 cells that incubated in various concentrations of Fulverstrant (D) or Letrozole (E) for 7 days. F, G. MCF-7 cells with/without MYSM1 knockdown were subjected to colony formation assay under diverse doses of Fulverstrant (F) or Letrozole (G). Clones were stained with R250 and photographed 2 weeks later.

---

**Supplementary Table 1.**
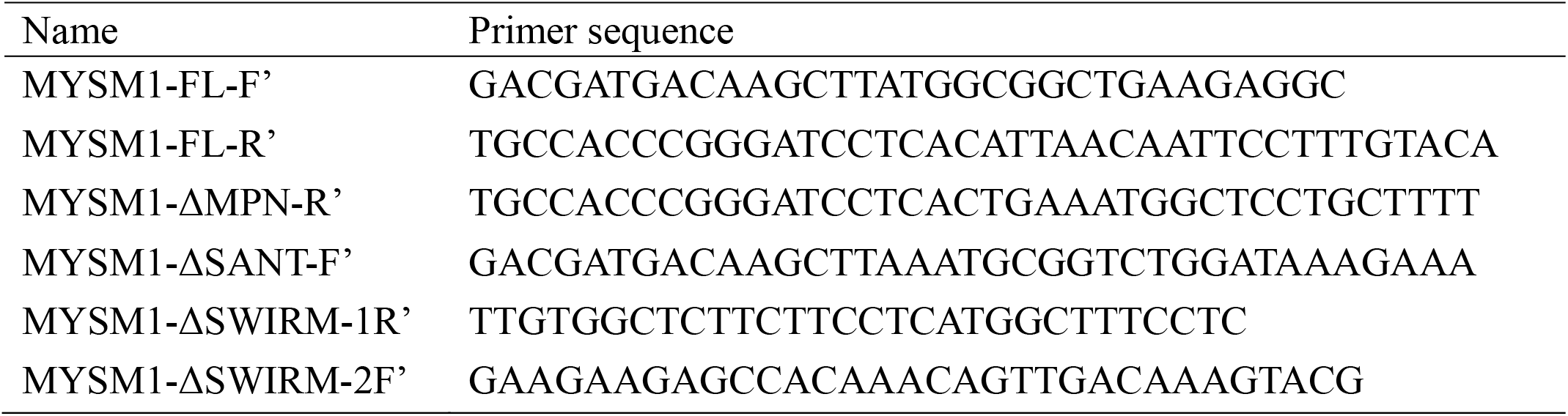
Primers used for MYSM1 plasmids construction in this study.

**Supplementary Table 2.**
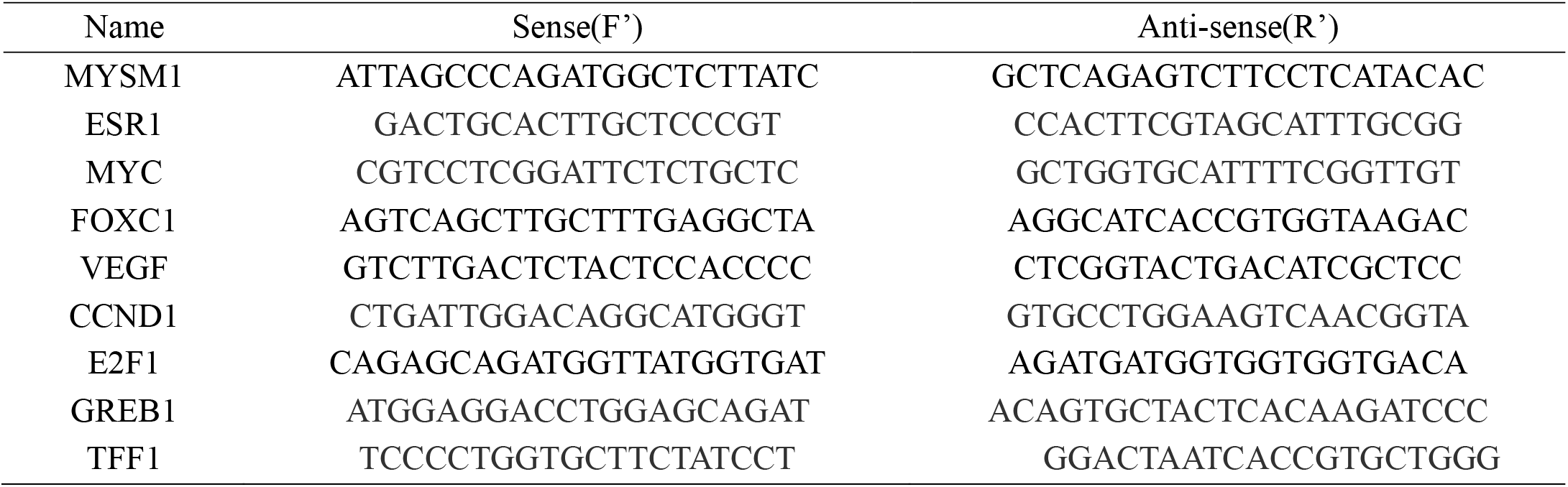
Primers used for qPCR.

**Supplementary Table 3.**
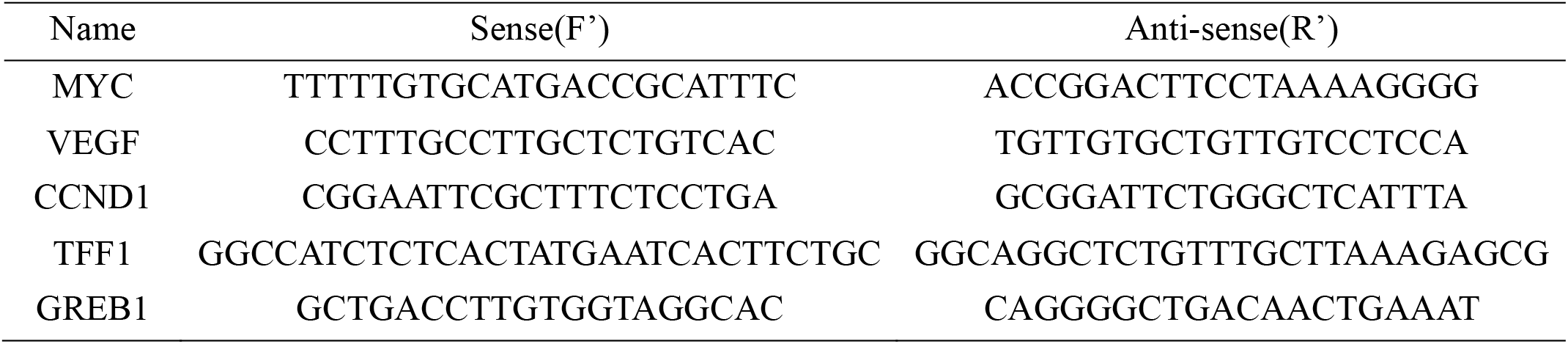
Primers of ChIP used for qPCR.

